# *UnifiedGreatMod*: A New Holistic Modeling Paradigm for Studying Biological Systems on a Complete and Harmonious Scale

**DOI:** 10.1101/2024.09.18.613635

**Authors:** Riccardo Aucello, Simone Pernice, Dora Tortarolo, Raffaele A. Calogero, Celia Herrera-Rincon, Giulia Ronchi, Stefano Geuna, Francesca Cordero, Pietro Lió, Marco Beccuti

**Author notes:** **Correspondence** Department of Computer Science, University of Turin, Via Pessinetto 12, 10123, Torino, Italy. Equally contributing authors (Riccardo Aucello, Simone Pernice). Equally supervised contribution (Francesca Cordero, Pietro Lió, Marco Beccuti). **Funding information**.

## Abstract

**Motivation:** Computational models are crucial for addressing critical questions about systems evolution and deciphering system connections. The pivotal feature of making this concept recognisable from the biological and clinical community is the possibility of quickly inspecting the whole system, bearing in mind the different granularity levels of its components. This holistic view of system behaviour expands the evolution study by identifying the heterogeneous behaviours applicable, for example, to the cancer evolution study.

**Results:** To address this aspect, we propose a new modelling paradigm, UnifiedGreatMod, which allows modellers to integrate fine-grained and coarse-grained biological information into a unique model. It allows for functional studies, in which the understanding of the system’s multi-level stable condition and the system’s fluctuating condition are combined to investigate the functional dependencies among the biological entities in the system under study. This is achieved thanks to the hybridisation of two analysis approaches that capture a system’s different granularity levels. The proposed paradigm was then implemented into the open-source, general modelling framework GreatMod, in which a graphical meta-formalism is exploited to simplify the model creation phase and R languages to define user-defined analysis workflows. The proposal’s effectiveness was demonstrated by mechanistically simulating the metabolic output of *Echerichia coli* under environmental nutrient perturbations and integrating a gene expression dataset. Additionally, the UnifiedGreatMod was used to examine the responses of luminal epithelial cells to *Clostridium difficile* infection.

## 1 INTRODUCTION

Traditionally, biological problems have been approached using a reductionist perspective by thoroughly dissecting biological phenomena into individual components. However, with the advent of deep sequencing technologies, there has been an unprecedented opportunity to comprehensively characterise biological components simultaneously and gain a global sense of the systemic interactions that occur within a cell (Feist *et al*., 2009).

In computational biology, the pursuit of understanding complex biological systems has led to the development of diverse modelling approaches, each offering unique rationales and challenges. Constraint- and mechanism-based modelling have emerged as powerful tools for interpreting biological phenomena. Specifically, constrain-based approaches, such as Flux Balance Analysis (FBA) (Machado and Herrgård, 2014; Opdam *et al*., 2017), can be employed to reconstruct genome-scale metabolic networks incorporating curated and systematised information about the known small metabolites and metabolic reactions of a cell type based on transcriptomics, proteomics, or metabolomics data (Bordbar *et al*., 2014). Furthermore, many studies have been trying to predict fluxes via transcriptomics and FBA or other constraint-based modeling techniques; recent approaches, such as (Damiani *et al*., 2019; Di Filippo *et al*., 2021; Galuzzi *et al*., 2022) map from expression profile to constraints for metabolic fluxes adjusting fluxes based on gene expression levels. Among the available tools for FBA the COBRA Toolbox is integral to the constraint-based modelling community, which focuses on open-source genome-scale metabolic models (GEMs). Noteworthy projects for high-throughput GEM generation include Path2Models (Büchel *et al*., 2013), AGORA (Magnusdottir *et al*., 2016), CarveMe (Machado *et al*., 2018), and BiGG Models (Norsigian *et al*., 2019). Moreover, human and microbial models and maps are available at the Virtual Metabolic Human website (Noronha *et al*., 2018), which collects human and gut metabolism data and links this information with nutrition. These modelling approaches are efficient for the analysis of large-scale metabolic networks, but need detailed descriptions of molecular mechanisms, such as quantitative information on molecular abundances and reaction kinetics. In contrast, fine-grained biological information is often modelled using systems of ordinary differential equations (ODEs), namely, mechanism-based approaches, to reproduce the dynamics of the system (Kendall *et al*., 1999). Although these approaches have the potential to reproduce reality more closely by investigating a biological phenomenon at a greater depth, they need help with scalability and parameterisation issues, as they require a considerable amount of preliminary quantitative information, which limits their applicability to larger systems.

Recognising the complementary nature of these approaches, there is growing interest in developing computational models that bridge the gap between constraint-based and mechanistic modelling paradigms. An attempt to expand the possibilities of FBA is Dynamic Flux Balance Analysis DFBA proposed in (Maranas and Zomorrodi, 2016; Mahadevan *et al*., 2002; Yang *et al*., 2019). DFBA represents a compromise between fully dynamic models - which cannot be simulated on a large scale - and steady state models - in which metabolite fluxes within a biological system remain constant over time and do not involve reaction kinetics (Palsson, 2011; Maranas and Zomorrodi, 2016). Its novelty lies in its ability to model metabolism under dynamic conditions, combining extracellular dynamics with intracellular steady states. However, while DFBA allows investigating genome-scale networks under transient conditions, it cannot implement mechanistic knowledge of non-metabolic processes.

Recently, a collective effort has been to hybridise mechanism-based and constraint-based approaches. The goal is to tackle their limitations and merge their capabilities to decrease costs, improve efficiency, and perform more descriptive phenotype predictions. A first insight was provided in (Pernice *et al*., 2020a), where the authors proposed some initial indications on how this hybridisation can be carried out through a high-level graphical formalism. In particular, the paper clearly shows how FBA can be exploited as a global source and sink of the system for specific sets of metabolites. Moreover, it discusses the possibility of using FBA in the network sub-models where there are missing kinetics and/or sloppy parameters, and to close the original system.

Another notable example was the recent proof of concept of a hybrid ODE- and constraint-based model proposed in (Ben Guebila and Thiele, 2021), where an ODE-based glucose-insulin model was combined with a whole-body organ-resolved reconstruction of human metabolism to investigate Type 1 diabetes mellitus. However, the computational methodology and associated framework used in such work are peculiar to the example proposed.

Thus, to the best of our knowledge, a well-formalised methodology that is sufficiently general is still missing. To address a such gap in this work, we advance the hybridisation of these two approaches by first formally defining the coupling between ODEs and FBA using a graphical meta-formalism, and second by implementing this new modelling paradigm, namely *UnifiedGreatMod*, into the well-known open source and general modelling framework *GreatMod* (Castagno *et al*., 2020) with the aim of provide general-purpose, scalable, reproducible, and easy-to-use modelling tool allowing researchers to study biological systems on a complete and harmonious scale.

Finally, it is important to emphasise that our framework is designed to work synergistically with the COBRA Toolbox, enabling users to seamlessly switch between or simultaneously utilise mechanistic-based and constraint-based models as needed. This interoperability ensures that users can effectively integrate existing COBRA models and tools with the models defined in *UnifiedGreatMod*, creating a cohesive and powerful modelling environment.

## 2 METHODS

The flow of the paradigm involves the integration of heterogeneous data, considering different biological domains of a cell (such as genomic, transcriptomic, and metabolic data) or as distinct entities interacting with each other, including microbes interacting with host cells, facilitating their interchange within a holistic view of biological systems.

The harmonisation, a central aspect of our paradigm, facilitates a dialogue among biological data and merges the computational aspects of the paradigm. Specifically, how the combination of mechanism and constraint-based models simulates the entire system to yield temporal evolution of key biological entities. This simulation enables the derivation of new insights into the systems under study.

*UnifiedGreatMod* aims to harmonise two different solution techniques: ODEs and FBA. This is achieved through a graphical formalism, which allows the modeller to state (i) the dynamic model represented by an ODEs system, (ii) the metabolic model studied by the FBA, and (iii) the coupling between the dynamic and metabolic models (see Fig.1).

### 2.1 Overview of the coupling definition

The innovative contribution of *UnifiedGreatMod* regards the harmonisation between the dynamic and metabolic models. Intuitively the key to this harmonisation is the identification of *metabolites* and *reactions* to be coordinated, assuming (i) the existence of a set of metabolites shared between the two models, representing the resources that might be produced or consumed by both models, and (ii) the existence of boundary reactions which allow the sharing of the respective metabolite between the models. The following describes the idea behind the harmonisation step, while all the theoretical details are provided in Supplementary Material S1.1.

**FIGURE 1.**
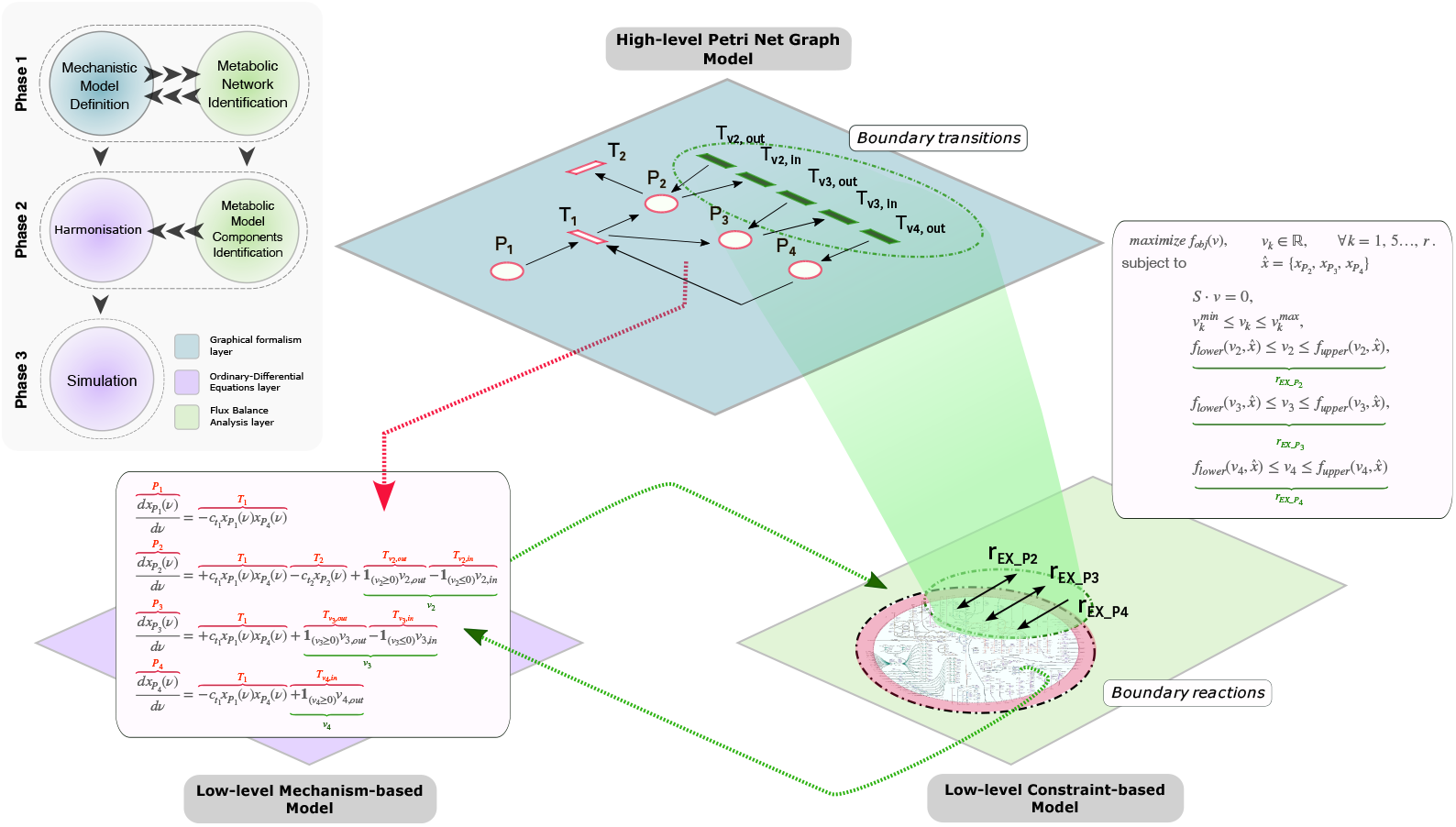
A sketch of the proposed modelling paradigm based on the graphical Petri Net formalism (i.e. ESPN) for stating the harmonisation between the mechanistic-based and the constrained-based models. The red colour is associated with the components of the ODEs derived from the ESPN, while the green one with the fluxes obtained from the FBA.

As shown in left-upper corner of Fig. 1, the *UnifiedGreatMod* workflow can be summarised in the following three phases:

1. definition of the mechanistic-based model and identification of constraint-based model;
2. identification of the reactions encoded in the constraint-based model (i.e., metabolic network) to exploit the harmonisation by coordinating the two modelling techniques;
3. simulation the unified system.

#### 2.1.1 Phase 1 - step 1: Definition of the mechanistic model

In this first step of the first phase, the user initially draws the mechanistic-based model using the graphical PN formalism. In detail, Petri Nets (PNs) (Marsan *et al*., 1995) and its generalisations, such as the Extended Stochastic PNs (ESPNs) (Pernice *et al*., 2020a), are a high-level mathematical formalism that exploits graphical elements to represent system components and their interactions. ESPN formalism has been extensively used to study biological systems (Pernice *et al*., 2023; Castagno *et al*., 2020; Pernice *et al*., 2020b) thanks to its ability to represent reaction systems in a natural graphical manner and provide qualitative and quantitative information on the system’s behaviour under study.

The basic components are (i) **places**, corresponding to state variables of the system (graphically represented as circles), and (ii) **transitions**, denoting events or activities (graphically represented as boxes). Places can contain *tokens* drawn as black dots (namely marking of the place), representing the modelled entities of the system. Then, the number of tokens in each place defines the state of a PN, called *marking*. Places and transitions are connected by *arcs*, which express the relation between states and event occurrences. Moreover, by definition, ESPN can successfully integrate complex velocities functions in a unique model, and combine metabolic and regulatory networks (see the *harmonisation* paragraph).

An example of ESPN is depicted in the first-top layer of Fig.1, which is characterised by four places (named **P1, P2, P3**, and **P4**) and seven transitions (named **T1, T2, T**_**v2**,**out**_ , **T**_**v2**,**in**_, **T**_**v3**,**out**_ , **T**_**v3**,**in**_, and **T**_**v4**,**out**_). For instance, the *T* 1 transition is connected to two input places (*P* 1 and *P* 4) and two output places (*P* 2 and *P* 3). Thus, it can change the marking of the four places through its *firing*, i.e., by removing tokens from *P* 1 and *P* 4, and adding tokens to *P* 2 and *P* 3 with a firing intensity defined by the Mass Action law: 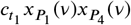. Where 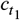 is the kinetic constant of the transition, and 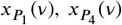 are the number of tokens at time *v* in the places *P*_1_ and *P*_4_, respectively.

As shown in (Pernice *et al*., 2020a), starting from an ESPN model, it is possible to derive the system of ODEs describing how the number of tokens changes over time in each place, as graphically represented by the red dashed arrow in Fig.1. For instance, if we consider the first ODE in the rectangle pointed by the red dashed arrow, its solution describes the evolution in the number of tokens in the place *P* 1 over time. Since *P* 1 has only one transition (*T* 1) that might change its, specifically by removing tokens with intensity defined as in the previous paragraph. The same for the remaining three places.

For the model construction and automatic derivation of the low-level mathematical processes characterising the system dynamics, we exploited and appropriately extended *GreatMod*, a general modelling framework to simulate biological complex systems using an intuitive graphical interface. *GreatMod* is composed of three main modules: (i) a Java GUI based on Java Swing Class, called GreatSPN (Amparore *et al*., 2016), which allows the user to draw models using PN formalism and its extensions, (ii) the R library *epimod* (Castagno *et al*., 2020), and (iii) Docker containerisation, a lightweight OS-level virtualisation. Further details are reported in the Supplementary Material S1.1.4.

#### 2.1.2 Phase 1 - step 2: Identification of the metabolic network

For metabolic network identification, the corresponding model can be encoded in MAT-file (“.mat”) format as defined by the COBRA Toolbox (Heirendt *et al*., 2017). Then, the constrained-based modelling approach, called Flux Balance Analysis (FBA), can be used to study metabolic networks by analysing the flow of metabolites.

A linear programming solver is thus used to find the flux distribution *v* ∈ ℝ^*r*^ of all the *r* reactions that maximise or minimise the objective function *f*_*obj*_ : ℝ^*r*^ → ℝ while satisfying two main constraints: (i) the steady-state assumption *S* · *v* = 0, and (ii) each flux must take value between 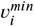 and 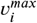, that are the lower and upper constraints of the *i*-th flux. Let us denote *F* , the set of all the fluxes *v*_*i*_, *i* = 1, … , *r*.

To facilitate the integration of a constrained-based model into our framework, we developed the “*epimod_FBAfunctions*” R package.This package offers essential functions to read and modify MAT files, translating metabolic network representations into the *GreatMod* internal format. In particular, using *epimod_FBAfunctions*, it is possible to integrate (i) gene expression data and (ii) dietary inputs to set the fluxes constraints properly. We have implemented a strategy to integrate metabolomics and gene expression data. Our computational pipeline can (i) compute differential reaction activities from transcriptomics to predict whether the differential expression of metabolic enzymes directly causes differences in metabolic fluxes and (ii) constrain boundary reactions based on the concentration of metabolites resulting from a given dietary input. The critical aspect of integrating transcriptomic data lies in selecting the integration method. The package *epimod_FBAfunctions* implements the integration process presented in (Kashaf *et al*., 2017). This method involves evaluating the Gene-Protein-Reaction rules associated with each reaction to derive reaction expression values from gene expression values.

#### 2.1.3 Phase 2 - step 1: Identification of the reactions

The first step of the second phase aims to identify a set of boundary reactions, also known as exchange or transport reactions, *F* ^*shared*^ ∈ *F* , so each corresponding flux can be associated with a velocity in the ODEs system, representing the exchange of such metabolites. These reactions define the input and output of metabolites to and from the metabolic network in FBA. These reactions are crucial for accurately modelling the system because they set limits on the availability and removal of metabolites. Firstly, we need to filter the set of all the fluxes of the FBA model to consider reactions that can be used to define *F* ^*shared*^ . Specifically, we strongly suggest using only the boundary reactions for two reasons: (i) to maintain consistency with the fundamentals of our paradigm and (ii) not to violate the balance assumption of the FBA model. Our new paradigm has to connect two modelling approaches with their assumptions and rules, simulating two different system layers which share a given number of resources (e.g., metabolites). In particular, the FBA model is constrained by the system’s steady state, which supposes no changes in any metabolite concentration. Therefore, by defining a single ODE that models the concentration over time of a metabolite also described in the FBA model, it is straightforward that the steady-state assumption of the FBA model does not hold anymore. Differently, the boundary reactions represent by definition the connection of the metabolic model with an extracellular environment, delineating in such a manner the perfect bridge between an ODE representing the shared metabolite outside the metabolic model, which acts as its resource by modifying dynamically the reaction constraints, respectively.

Among the boundary reactions, the ones with a greater sensibility to perturbations in their constraints play a central role in connecting the two modelling approaches. Given that the system of ODEs is connected to the FBA model in terms of the fluxes constraints, it is straightforward to consider reactions where small perturbations in the concentration of the shared metabolite affect the FBA solution. Otherwise, the two approaches could be solved independently without dynamic communication, which might improve and change the analysis. Therefore, we suggest focusing the modelling efforts on high-sensitive reaction fluxes to minimise the poor call of such boundaries and reduce the likelihood of errors. We focus on fluxes for which a small perturbation entails a substantial variation of the FBA model outcomes (e.g., the objective function value). Specifically, by exploiting Sobol’s variance-based Sensitivity Analysis (SA) (Saltelli *et al*., 2010), a powerful and widely-used method to determine the contribution of input variables to the output variability of a mathematical model, it is possible to detect automatically critical fluxes in genome-scale metabolic, which is particularly useful for understanding how changes in the fluxes constraints influence the objective function. In this context, we developed in *epimod_FBAfunctions* package functionality to perform the SA considering a constrained-based model.

Further details regarding the definition of *F* ^*shared*^ and the definition of the SA approach are reported in the Supplementary Material S1.3 and S1.1.3, respectively.

#### 2.1.4 Phase 2 - step 2: Harmonisation

Once *F* ^*shared*^ is defined, the next step is defining how to coordinate the two modelling approaches. The general idea is that the concentrations obtained from solving the ODEs could be used as flux constraints because they represent the availability of the metabolites in the environment and, therefore, the uptake limit. Usually, the upper constraint is set only if a growth limit has to be defined. Otherwise, it should be infinity (see the Supplementary Material S1.4).

Thus, given an ESPN model, we consider *T*_*g*_ the set of all transitions (called general transitions) whose firing intensities are defined as continuous real functions, and not through the Mass Action law. Let us refine *T*_*g*_ into two disjoint subsets 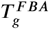 and 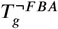 , so that i) 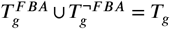, and ii) 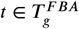 iff its intensity is obtained by solving the constrained-based model. Let us observe that, by definition, the transition intensity has to be a positive number, therefore if reversible reactions are considered in *F* ^*shared*^ , then they are decoupled into irreversible reaction pairs, defining two different transitions, denoted with _*in* if the estimated flux is negative and _*out* if it is positive. Thus, we can infer that

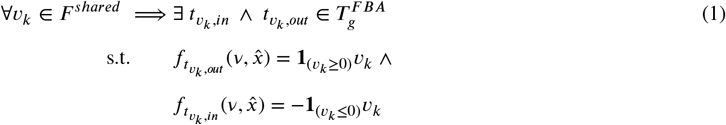

where, *f*_*t*_() is the firing intensity associated with the general transition *t* depending on the marking of its inputs places 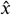 at time *v*, 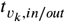 the general transition representing the flux *v*_*k*_, and **1**_(·)_ is the indicator function that returns 1 only if the input condition is satisfied.

As depicted with the green colour in Fig.1, *F* ^*shared*^ is defined by three fluxes *v*_2_, *v*_3_, *v*_4_ associated respectively with the boundary reactions 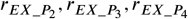. Let us observe that these three reactions were selected since they represent the exchange of the metabolites represented by the places *P*_2_, *P*_3_, and *P*_4_, respectively. Therefore, for each reaction, one or two general transitions are defined depending on whether it is a reversible reaction or not. For instance, the reaction 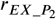 is associated with two general transitions 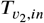 and 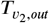, whose intensity will depend on the sign of *v*_2_ obtained by solving the FBA.

Let us observe that the constraints of the shared fluxes are defined as functions 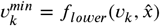 and 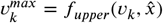 (lower and upper bounds, respectively) that depend on the solution of the system of ODEs thought 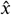, i.e., the input places of the transitions in 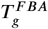. For instance, they might be defined as a linear transformation of the respective exchanged metabolites concentration (i.e.,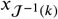) when *i* ∈ *M*^*Ex*^.

#### 2.1.5 Phase 3: Simulation

Finally, the simulation technique integrates the solution of the system of ODEs, which is based on the Backward Differentiation Formula method (Burden *et al*., 2016), with the resolution of the Linear Programming Problem^1^. Specifically, if at least one of the metabolite concentrations is modified during the solution of the system of ODEs, that is, some lower and upper bounds might change, then the LPP is solved with the new constraints, and the fluxes used in the ODEs are updated (see the Supplementary Material S1.5).

An example of the final system of ODEs is depicted in Fig.1, in which both the components of the mechanistic-based (in red) and constrained-based (in green) models are present.

## 3 RESULTS

The use of deep sequencing technology to analyse microbiome profiles across different tissues has prompted the microbiology community to expand their research objectives to include the functional aspects of microorganisms integrating heterogeneous data. For this reason, we propose two applications of *UnifiedGreatMod* in this context.

The first application is related to the carbon catabolite repression phenomenon exhibited by *E. coli* (Ammar *et al*., 2018), with all details provided in the Supplementary Materials. The second application, described hereafter, investigates the interaction between *Clostridioides difficile* and intestinal epithelial cells (IECs).

### Definition of mechanistic and constraint-based models

*C. difficile* is a gram-positive bacteria that causes infections typically affecting older adults in hospitals or long-term care. Symptoms of the clinical disease can range from diarrhoea to life-threatening damage to the colon. During infection, *C. difficile* colonises the large intestine and causes toxin-mediated disruption of the intestinal epithelial barrier function, which results in increased intestinal permeability and triggers haemorrhage (Kouhsari *et al*., 2018). Recently, evidence has shown that chronic *C. difficile* infection contributes to colorectal tumorigenesis (Drewes *et al*., 2022). To model the interaction between *C. difficile* and IECs during infection, six interacting biological domains have been considered: (i) *C. difficile metabolic network* incorporates all metabolic reactions of the *C. difficile*; (ii) *C. difficile cell dynamics* models the proliferation response of a bacteria population upon treatment with antimicrobial therapy; (iii) *metronidazole action* describes the mechanism of action of a drug driving an imbalance between oxidative and antioxidative processes, inducing oxidative stress and cell death; (iv) *intestinal lumen* models the toxin-mediated inflammation and the dietary intake of nutrients; (v) *intestinal epithelial cells* describes the dynamics of the IEC and the absorption of nutrients, and (vi) *blood vessels* describes the transepithelial amino acids transported into the circulation following protein digestion and absorption. The detailed biological description of all modules is provided in Supplementary Material S3.1. The model’s parameterisation is primarily based on values extracted from the literature, except for *Death4Treat, Detox*, and *IECsDeath*. These parameters are explored using Partial Rank Correlation Coefficients (PRCC) (Fornari *et al*., 2015) to observe changes in the overall model dynamics (see Supplementary Figure S5). Further details are available in Supplementary Material S1.2.1. Additionally, Supplementary Material S2 provides a detailed example of how parameters can be derived from transcriptomic data.

### Model component identification and model harmonisation

Harmonising the low-level ordinary differential equations and the low-level metabolic model (exchanges between the external environment and the metabolic model) involves identifying the critical fluxes through Sobol’s variance-based Sensitivity Analysis (SA) (Saltelli *et al*., 2010). The SA method indicates that *EX_leu_L_e* and *EX_trp_L_e* are the two transitions in our model with the highest influence on the output. To better capture all dynamics biologically related to the antibiotic-resistant mechanism (Karasawa *et al*., 1995; Knippel *et al*., 2020), we also consider the transitions *EX_pro_L_e, EX_ile_L_e, EX_val_L_e, EX_cys_L_e*, and *sink_pheme_c*. These reactions are split into *in* and *out* components to retain direction information.

### Modelling antibiotic therapy across experimental settings and conditions

The coupled model runs simulations to predict the system behaviour under different experimental settings and conditions. To assess the significance of *UnifiedGreatMod*, we proposed three experiment settings. In the first, called *ablation experiment*, the dynamics of the metabolic environment are computed by FBA only at the beginning of the simulation. In the second experiment, named *partial-ablation experiment*, the FBA is computed when an external stimulus occurs, i.e., at any drug injection. In the last experiment, namely *unified experiment*, the FBA model is continuously solved for the state of the ODEs system (Fig. 2A). Moreover, system dynamics are also responsive according to different systems conditions, where a condition is represented by given systems parameters configuration selected after a calibration step aimed to i) minimise the IECs damage, ii) keep the drug concentration closed to the minimum inhibitory concentration value, and iii) minimise the number of *C. difficile* cells without complete eradication. As expected, in the unified setting, in a condition of limited drug efficacy, the successful colonisation by *C. difficile* leads to an increment in the number of *C. difficile* cells up to a plateau value. Consequently, IECs die, the gut lining is damaged, and, following extravasation, red blood cells undergo lysis, which increases heme concentration in the gut microenvironment Fig. 2B. The drug therapy is modelled as an injection at specific time steps (i.e., 8, 16, 24, 32, 40, and 48 hours). In the Unified scenario, it is possible to note a reduction in the number of *C. difficile* cells at the time of the drug injection, while the number of IECs decreases since the deadly activity of the infection, the intracellular heme concentration increases. An overview of the model results for all places is reported in Supplementary Fig. S4.

**FIGURE 2.**
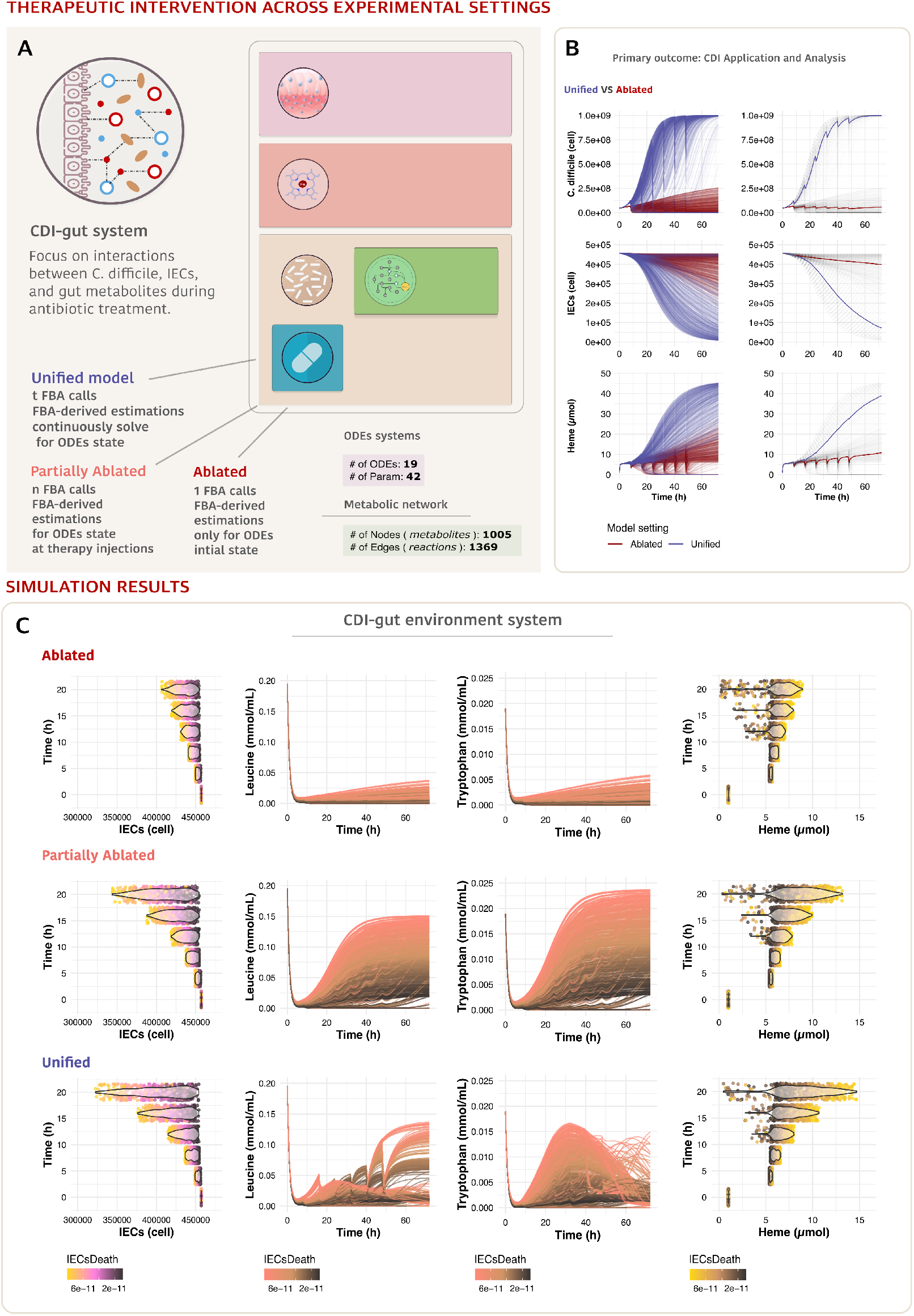
*C. difficile* infection model. (A) the model designed scheme, (B) *C. difficile*, IECs number and intracellular heme concentration coloured by experimental settings, (C) IECs number, leucine tryptophan, and intracellular heme concentrations coloured by different virulence conditions (gradient of *IECsDeath* transition values) for the three experimental settings.

### Simulation results: comparison of ablation versus *UnifiedGreatMod* models

The comparison among the three experimental settings shows different trajectories during the system’s evolution Fig. 2C. For example, in all three settings, the number of IEC cells changes linearly with the *IECsDeath* rate (which represents the degree of infection virulence). However, the effects of the unified model simulations become evident when observing the number of cells: specifically, increasing cell death leads to a significant decrease in cell numbers in the unified model compared to the ablated experimental settings. Other distinct behaviours are prominent in each experimental setting when examining the trajectories of leucine, tryptophan and intracellular heme (Fig. 2C). In the unified experiment compared to ablated and partially ablated configurations, lower values of the *IECsDeath* parameter are sufficient to decrease the IEC number resulting in a differential perturbed metabolic gut microenvironment.

Furthermore, the unified experiment is the only setting able to unveil the presence of a heme-mediated antibiotic-resistant strains spectrum (from fully susceptible, to partially susceptible and resistant populations), where the model’s results illustrate how a gradient of conditions enables model richer collection of antibiotic resistance types. (Fig. 2B).

## 4 DISCUSSION

Computational modelling is becoming increasingly crucial to formulate new hypotheses and provide plausible explanations of how a real system works. In-silico models may help uncover how system components interact with each other and how they are related to wider biological effects. However, new computational theoretical and practical achievements are needed to model how cell dynamics are intertwined with the mechanics of cell signalling, metabolism, and the influence of extracellular factors. Here, we propose *UnifiedGreatMod* based on open source and general framework to integrate formal concepts and different solution techniques and offer the possibility of relating multi-level dynamics. *UnifiedGreatMod* exploits a single easy-to-use graphical meta-formalism ensuring result reproducibility. The most innovative aspect of *UnifiedGreatMod* is the possibility to continuously compute the evolution of the systems considering at the same time, all the reactions involved in the metabolism, and changes in the micro-environment. Due to its clinical importance as a nosocomial disease and an emerging tumorigenic pathogen, *UnifiedGreatMod* was exploited to model *C. difficile* infection in the human gut as a case study.

In the gut, the growth medium composition significantly affects *C. difficile*’s toxin production. These toxins compromise the epithelial cell barrier, causing apoptosis in IECs. Toxin synthesis regulation is a complex bacterial response to specific nutrient availability during infection. *C. difficile* virulence is tightly linked to nutrient availability; when nutrients are limited, the bacteria produce toxins. These toxins release host-derived nutrients through cytotoxic activity. When IECs die, they release their intracellular contents, including nutrients such as amino acids, into the gut environment. These nutrients then become available for the bacteria.

Leucine is a key metabolite of *C. difficile* Stickland fermentation, i.e., it can be used to generate ATP through its paired oxidation and reduction with other amino acids. It has been shown that leucine is one of the main energy sources during bacterial exponential growth and that its pools are depleted before the onset of the stationary phase (Hofmann *et al*., 2018). The role of tryptophan in the growth of *C. difficile* is not explicitly defined in the literature. However, *C. difficile* can not synthesise tryptophan, as suggested by (Neumann-Schaal *et al*., 2019). It would need to obtain this amino acid from the environment, which could influence its growth dynamics.

The comparison among ablated, partially ablated, and unified experiments demonstrated that simulations depict the connection between nutrient availability in the gut and bacterial growth state during the disease. In all scenarios, tryptophan and leucine dynamics are associated with the model parameter representing the cytotoxic capacity of the bacterium upon IECs. This parameter reflects the degree of toxic gene regulation by nutrient-sensing transcriptional repressors, which are depressed with low levels of environmental nutrients (Theriot and Fletcher, 2019)

Nevertheless, by dynamically computing the FBA throughout the entire simulation, we can uncover mechanisms of antibiotic resistance. This mechanism may promote chronic infection by *C. difficile*, even when metronidazole treatment is ongoing. Antibiotic injections reduce the number of bacteria at drug administration. Ablation experiments simply indicate that infection clearance reduces damage to IECs. In contrast, the latest experiments better simulate the system’s behaviour by representing a greater richness, showing *C. difficile* entering two nutrient-dependent stationary phases upon drug treatment. The two states are visible from the *C. difficile* cells dynamics. Highly susceptible bacteria are cleared, preserving IEC barrier integrity and reducing nutrient leakage, while highly resistant bacteria kill IECs, saturating the environment with nutrients. Intermediate susceptibilities are also depicted, reflecting real bacterial populations, which typically display a range of antibiotic susceptibilities rather than discrete categories. This gradient allows the model to show how intermediate types are selected and how to establish antibiotic resistance. Additionally, the two FBA integrated experiments capture the dynamic consumption of metabolites due to oscillations in the bacterial population with varying detail.

In conclusion, these results indicate that this new paradigm, harmonising dynamic and metabolic models, can significantly enhance personalised healthcare in the context of dynamic treatment and patient monitoring. ODEs allow for the precise modelling of temporal changes in biological systems, capturing the dynamic behaviour of disease progression and treatment response. Meanwhile, FBA enables the optimisation of metabolic pathways by analysing the flow of metabolites through networks, ensuring that cellular processes are efficiently managed. By integrating these two powerful approaches, healthcare providers can develop highly tailored therapeutic strategies that are continuously adjusted based on real-time patient data.

## Availability

GreatMod https://qbioturin.github.io/epimod/, epimod_FBAfunctions https://github.com/qBioTurin/epimod_FBAfunctions, first case study E.coli https://github.com/qBioTurin/Ec_coli_modelling, second case study C.difficile https://github.com/qBioTurin/EpiCell_CDifficile.

## Acknowledgements

This project has received funding from: Ministero dell’Univerisita’ e della Ricerca (MUR) PRIN 2022 project *MEDICA: Modelling and vErification of alkaptonuria and multiple sclerosis Driven by biomedICAl data* [No 2022RNTYWZ] (MEDICA project to Marco Beccuti) and from the European Union’s Horizon 2020 research and innovation program under grant agreement no. 825410 (ONCOBIOME project to Francesca Cordero and Simone Pernice) The scientific activities of the CINI InfoLife

Laboratory supported this research.

## Supplementary Material

### S1.1 Theoretical Background

#### S1.1.1 Petri Net formalism and its extensions

Petri Nets (PNs) Marsan *et al*. (1995) and their extensions are widely recognised to be a powerful tool for modelling and studying complex systems thanks to their ability to represent systems in a natural graphical manner and of allowing the computation of qualitative and quantitative information about the behaviour of these systems. In the literature, several generalisations of this formalism are presented to expand the possibility of modelling complex systems of different kinds (Herajy *et al*., 2018; Pernice *et al*., 2019). In this work, we will exploit the Extended Stochastic PN (ESPN), a PN generalisation allowing the definition of complex rate functions.

More specifically, ESPNs are bipartite directed graphs with two types of nodes: *places*, which correspond to state variables of the system, graphically represented as circles, and *transitions*, which correspond to the events that can generate a state change and graphically are represented as boxes. Places can contain *tokens* drawn as black dots, representing the modelled entities of the system. Then, the number of tokens in each place defines the state of an ESPN, called *marking*.

Nodes of different types are connected by *arcs*, which express the relation between states and event occurrences. A specific multiplicity is associated with each arc, and it describes the number of tokens removed from (or added to) the corresponding place upon the firing of the transition to which the arc is connected. Graphically, it is written beside the arc, but the default value of one is omitted. Functions *I* and *O* describe transitions’ input and output arcs, respectively. For convenience, these can be represented by *n*_*t*_ × *n*_*p*_ matrices of natural numbers, where *n*_*t*_ is the size of the transitions set *T* , and *n*_*p*_ of the set of places *P* . The matrix *L* = *O* − *I* is called the *incidence matrix* (Colom and Silva, 2006), representing the transitions’ overall effect.

Finally, the evolution of the system is given by the firing of enabled transitions, where a transition is enabled if each input place contains several tokens greater than or equal to a given threshold defined by the multiplicity of the corresponding input arc. Enabled transitions may *fire*, removing a fixed number of tokens from its input places and adding a fixed number of tokens into its output places (according to the multiplicity of its input/output arcs).

Finally, each transition is associated with a specific intensity, representing the parameter of the exponential distribution that characterises its firing time. In this context, to easily model events with velocities defined by different and complex functions, the transitions are split into two subsets: *T*_*ma*_ and *T*_*g*_ . The former subset contains all transitions that fire with a velocity expressed in Mass Action (MA) law (Voit *et al*., 2015). The latter includes all transitions whose random firing times are defined as continuous real functions. Hence, we will refer to the transitions belonging to *T*_*ma*_ as standard transitions and as general transitions those in *T*_*g*_ .

Let define ^**·**^**t** the subset of *P* containing the input places to transition *t*, and 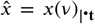 as the subset of the marking *x*(*v*) concerning just the input places to transition *t*. Thus, given a transition *t* ∈ *T* = *T*_*ma*_ ∪ *T*_*g*_ at the time *v*, it will move tokens in state *x*_*i*_(*v*) with speed defined as follows: (i) 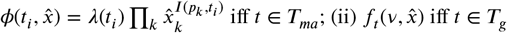. Thus, the instantaneous changes of tokens *x*_*i*_(*v*) in the *i*-th place at time *v*, is modelled by the following ODE:

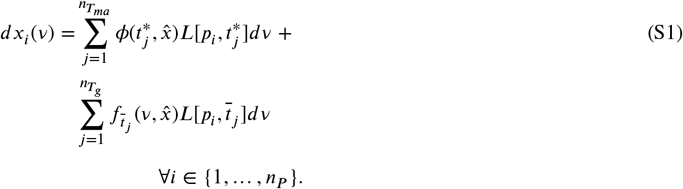

#### ESPN formal definition

##### Extended Stochastic Petri Net

An Extended Stochastic Petri Net (ESPN) system is a tuple (*P* , *T* , *I, O*, **m**_0_, λ, Λ), where:

- *P* = {*p*_*i*_} is a finite and non empty set of *places*, with *i* = 0, … , *n*_*p*_, where *n*_*p*_ is the number of places.
- *T* = *T*_*ma*_ ∪ *T*_*g*_ is a finite, non-empty set of transitions, with 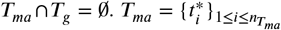 is the set of the 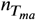 transitions whose speeds follow the MA law. 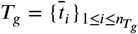 is the set of the 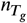 transitions whose speeds are defined as continuous functions.
- *I, O* : *P* × *T* → IN are the *input, output*, that defines the net’s arcs and specifies their multiplicities.
- **m**_0_ : *P* → IN is a multiset on *P* representing the *initial marking* of the net.
- λ : *T*_*ma*_ → ℝ gives the firing intensity of the transitions.
- Λ = {*f*_1_, … , *f*_*h*_} is the firing intensity set grouping the functions characterising the general transitions in *T*_*g*_ , with 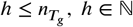. In particular, a function *f* depends only on the marking of the input places of the respective transition *t* (|^**·**^**t**|) and on time *v* ∈ ℝ^+^, i.e.,

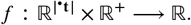

For instance the function *f*_1_ might represent a Michaelis Menten kinetic and *f*_2_ an Hill kinetic.

#### S1.1.2 Flux Balance analysis

Flux Balance Analysis (FBA) is a computational approach used in Systems Biology to study the metabolism of biological systems, particularly in microorganisms like bacteria and yeast (Papoutsakis, 2000; Watson, 1984). FBA is a powerful computational tool for predicting cellular metabolism by formulating it as a linear optimisation problem, where the goal is to find the optimal distribution of metabolic fluxes within a cell to achieve a specific biological objective, by assuming the steady state of the system. The first works ((Papoutsakis, 2000; Watson, 1984)) regarding the FBA date back to the early 1980s and showed the possibility of deriving from a system of metabolic reactions the stoichiometric equations describing the relations among different products and biomass and how to exploit linear programming for deriving the fluxes of such relations. FBA models have wide applications for simulating genome-scale reconstructions of metabolic networks. These networks are distinguished by a multitude of reactions and metabolites. The model operates under the assumption of a system-wide steady state. In particular, the flux distribution within the model is estimated by setting lower and upper boundaries for each flux. This function acts as a surrogate, representing the most plausible physiological state among all potential states of the system.

From a mathematical point of view, cellular metabolism is represented as a stoichiometric matrix that describes the relationships between different metabolites and reactions in the network. Each row of the matrix corresponds to a metabolite, and each column corresponds to a reaction. Fluxes represent the rate of flow of metabolites through each reaction in the network. FBA assumes that the cellular system is at a steady state, meaning that the concentrations of metabolites do not change over time, supposing that there is no net surplus or deficit of any metabolite. Mathematically the FBA modelling *m* metabolites and *r* reactions can be translated as a linear programming problem (LPP) as follows:

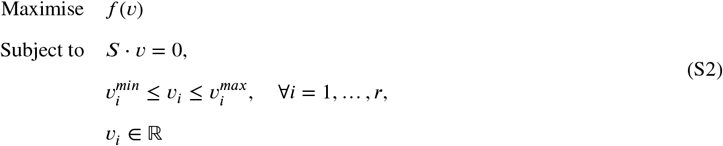

with *v* ∈ ℝ^*r*^ is the flux vector describing the activity of all the *r* reactions, *S* ∈ ℤ^*r*×*m*^ is the stochiometric matrix, *f* : ℝ^*m*^ → ℝ is the objective function to maximise, 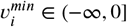 and 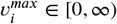 are the constraints of the *i*-th flux.

Constraints are applied to limit the allowable flux through each reaction, often based on experimentally determined rates or other biological constraints, and they are set depending on the type of the reaction ((Hao *et al*., 2010)): *exchanges* (uptake or secretion of metabolites into or out of the system), *transporters* (movement of metabolites across cellular membranes or compartments within the cell), *internals* (biochemical transformations that occur within the cell), and *demands/sinks* (need for specific metabolites within the system, or removal of certain metabolites without specifying a source).

In particular, these constraints are set depending on the type of the reaction. Typically metabolic models might be defined by more than one compartment (differential localisation of biochemical reactions within the cell), and so different types of reactions such as *exchanges, transporters, internals*, and *demands/sinks* can be defined depending on how they communicate within or among the compartments. The objective function is defined as a function *f* : *E* ⟶ ℝ where *E* represents the flux vector satisfying (i) the mass balance equation, and (ii) the constraints (both defined in Eq.s S2).

*Exchanges* and *demands/sinks* reactions define the different types of predefined *boundary reactions* because they define the limits or boundaries of the metabolic system being modelled. The term *boundary reactions* refers to reactions that involve metabolites entering or leaving the system, however, *exchanges* are usually connected to the extracellular compartment while *demands/sinks*, on the other hand, are typically connected to intracellular compartments.

*Exchange* reactions represent the uptake or secretion of metabolites between the cell and its surrounding environment. If an exchange reaction is only an uptake (import) reaction, then 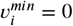 and 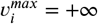. If it is only a secretion (export) reaction, then 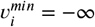 and 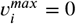. If the exchange reaction can function in both directions (import and export), then 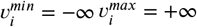. For a demand reaction (which consumes a metabolite), 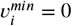 and 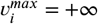. For a sink reaction (which can either consume or produce a metabolite) 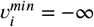 and 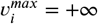.

Conversely, *transporters* and*internals* are types of predefined *core reactions* because they refer to reactions that involve metabolites entering or leaving the system’s compartments. *Transporters* represent the transfer of metabolites between two different compartments within the cell. The directionality and limits of these reactions can vary depending on the specific biological system being modelled. *Internal* reactions represent the conversion of one substance into another within the same compartment. Typically, for irreversible internal reactions, 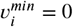 and 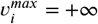. For reversible internal reactions, 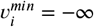 and 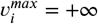.

#### S1.1.3 Global Sensitivity Analysis

FBA is a potent computational method for analysing metabolite flow through a metabolic network. However, the accuracy of FBA predictions is strongly influenced by the choice of flux boundaries, which define the feasible solution space that satisfies the system’s constraints.

Inaccurate flux boundary selection can lead to potential errors or unrealistic predictions that do not accurately represent the system’s actual behaviour. This issue is particularly critical when choosing flux boundaries for reactions that introduce nutrients into the system, as these boundaries play a crucial role in ensuring biologically relevant and accurate predictions.

To mitigate the risk of such errors, we propose focusing modelling efforts on high-sensitive parameters. These are parameters where minor perturbations can lead to significant variations in model outcomes. By concentrating on these parameters, we can minimise the impact of poor boundary selection and reduce the likelihood of errors.

By integrating global Sensitivity Analysis with FBA, we can enhance the reliability of metabolic models and provide more accurate predictions of metabolite flow through the network.

##### SA formal definition

In our study, we utilised Global SA as a computational strategy to examine the impact of various input uncertainties on the output of the FBA model. This approach has been widely used in previous studies to examine complex models, typically presenting sensitivity coefficients for model inputs in sorted order ((Qian and Mahdi, 2020)).

Our primary objective was to investigate how uncertainties in reaction boundary values influence the objective function of the FBA model. This function signifies the fluxes anticipated to affect the optimal cellular growth rate, which is characterised as the maximisation of biomass production.

We considered scenarios where the indicative value of each parameter was absent. It is a common practice in genome-wide modelling to assign a negative sign to intake fluxes, indicating a direction from outside to inside. To quantify the impact of parameter perturbation corresponding to the lowest boundary, we performed a variance decomposition in the domain [−10, 0] (mmol/gDW/h). We evaluated uncertainties by confining constraints within this interval, based on practical considerations and biological relevance. This interval has been shown to yield biologically relevant results and allows for a broad range of possible flux values ((Lularevic *et al*., 2019)).

Also, this interval disposition is a practical method to model nutrient influx boundaries. This approach is designed to establish boundaries that are low enough to accommodate internal reactions and high enough to include the stoichiometric coefficients of reactions involving these nutrients. The aim is to prevent internal boundaries from becoming limiting and to globally sample all possible ratios between intake fluxes by varying all parameters simultaneously, rather than one at a time.

Given the set of variables and with the intervals of variation of each input, an SA was executed into the following analytic steps: (i) the testing model parameter configurations were sampled, (ii) the model output was evaluated for each parameterisation, (iii) finally model output is collected and used to compute sensitivity coefficients. FBA optimisations are mutually independent, hence (ii) was performed in parallel. The fbar R package provided a toolkit for FBA and related metabolic modelling techniques. We performed Sobol’s variance-based SA relying on the *sensobol* R package ((Puy *et al*., 2022)), which provides implementations to conduct variance-based uncertainty and sensitivity analysis. Given D parameters in the model, the function *sobol_matrices()* allows for *N* * (2 * *D* + 2) independent random samples of the parameters space, where *N* is a user-defined parameter.

The method determines two kinds of sensitivity indices. The first-order indices *S*_*i* measuring the contribution to the output variance by a single model input alone:

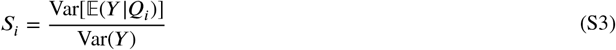

where *S*_*i*_ is calculated according to the sensitivity of a single parameter *Q*_*i*_ on the output variance of the model. The numerator represents the variance of the expected value of the output *Y* given the parameter *Q*_*i*_, while the denominator represents the variance of the output *Y* . The result represents the fractional contribution of the parameter *Q*_*i*_ to the output variance of the model. The total-order index *S*_*Ti*_ measures the contribution to the output variance caused by a model input, including both its first-order effects and all higher-order interactions:

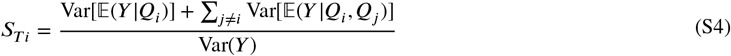

where the sensitivity of a single parameter *Q*_*i*_ on the output variance of the model, including the sensitivity due to interactions (covariance) between the parameter *Q*_*i*_ and all other parameters. Both the variance of the expected value of the output *Y* given the parameter *Q*_*i*_ and the variance of the expected value of the output *Y* given the parameters *Q*_*i*_ and *Q*_*j*_; where *Q*_*j*_ represents all other parameters except *Q*_*i*_.

##### SA implementation

In our study, we implemented global SA to classify reactions in an FBA model based on their impact on model outcomes. This method, although computationally intensive, allows for an extensive exploration of input factors. We utilised Sobol’s variance-based SA, combined with a random sampling scheme of parameters, to estimate the sensitivity indices of input parameters (Saltelli *et al*., 2010).

Our approach involved generating a set of samples using Sobol’s Quasi Random Numbers sequences, which were then used to compute the model’s output for each sample (Saltelli *et al*., 2010). We applied this method to the clostridioides reconstruction Clostridium difficile CD196, which required altering 186 boundary reactions and performing about 1.5 million FBA optimisations. We relied on the VMs+Storage services offered by the High-Performance Computing for Artificial Intelligence (HPC4AI) laboratory at the University of Turin. Due to the mutual independence of all linear programming optimisations, we offload computation to a parallel architecture with more than 5k cores. We leveraged the independence of FBA optimisations by distributing them across multiple cores where the parameter values were sampled using a uniform distribution, and the sample matrix was divided into chunks, each containing 12000 parameter configurations.

##### SA application

Genome-scale metabolic networks encapsulate all enzymatic reactions encoded in a given genome. In our study, we focused on the metabolism of *C. difficile*, which comprises over 1300 biochemical reactions whose analysis remains challenging due to largely restrictive and undetermined information, such as kinetic parameters of rate laws and the concentration of chemical species.

In the Clostridium difficile CD196 metabolic network, there are a large number of reactions (186) that exchange with the model’s external boundary. Estimating the value of each extracellular flux may not be feasible due to this large number. Therefore, it’s crucial to quantify the influence of these constraints on the model outputs to determine the most sensitive inputs. In our specific scenario, this approach facilitated the exploration of the factors involved in the cross-talk between different components of the system, represented by varying levels of analytical detail.

We downloaded the metabolic reconstruction of Clostridium difficile CD196 from the GitHub repository hosting AGORA (Assembly of Gut Organisms through Reconstruction and Analysis). The repository contains the Clostridium difficile CD196 metabolic model for AGORA version 1.03 in MATLAB format. We manually edited the model by incorporating boundary reactions not included in the reference reconstruction. The reversible sink reaction for heme:

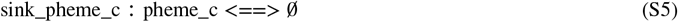

was added to enable the simulation of metabolic behaviour under changes in heme availability or genetic perturbations. By defining the heme inputs and outputs of the system, the heme sink reactions provide a framework for integrating heme dynamical detail into the model. Such an augmented model contains 1368 reactions (including 186 boundary reactions), 1005 metabolites, and 889 genes. We investigated the sensitivity of all the D=186 boundary reactions that allow the exchange of metabolites between the system being modelled. In this regard, we defined the parameter N equal to 2^1^3, resulting in *N* · (*D* + 2) = 1540096 independent optimisations distributed across 24 cores. This global SA pipeline ranks boundary reactions based on sensitivity, aiding in the identification of metabolites and reactions sensitive to the objective function of an FBA model, specifically biomass maximisation.

#### S1.1.4 GreatMod framework

*GreatMod* (https://qbioturin.github.io/epimod/) is a general modelling framework to simulate biological complex systems using an intuitive graphical interface for model construction and automatic derivation of the low-level mathematical processes characterising the system dynamics.

GreatMod is composed of three main modules. The first consists of a Java GUI based on Java Swing Class, called GreatSPN (Amparore *et al*., 2016), which allows the user to draw models using a high-level graphical formalism called Petri Net (PN) and its extensions. The other two modules implement all the functionalities needed for the model analysis: (i) the R library *Epimod* (Castagno *et al*., 2020) and (ii) Docker containerisation, a lightweight OS-level virtualisation.

Epimod provides a user-friendly interface for easy access to analysis techniques for (i) the derivation of the model’s underlying mathematical processes (e.g., the system of ODEs), (ii) the model sensitivity analysis to reduce the search space associated with each unknown parameter, (iii) the model calibration analysis to adjust the parameters to obtain the expected behaviour with respect to the available data, and (iv) the model simulation to answer specific questions through what-if analysis and to derive new insights. Finally, the Docker containerisation of all implemented analysis techniques ensures results reproducibility and framework portability, as specified by the RBP guidelines (Kulkarni *et al*., 2018).

### S1.2 The Harmonisation Paradigm

#### S1.2.1 Harmonisation definition

The key to harmonising dynamic and metabolic models lies in identifying metabolites and reactions to be coordinated. This involves assuming (i) shared metabolites between the models, which represent resources produced or consumed by both, and (ii) boundary reactions that facilitate the sharing of these metabolites. Our goal is to minimise essential parameters requiring measurement and to pinpoint influential parameters crucial for harmonisation. By integrating global SA with ODE-based models, we address the non-linear relationships between the most sensitive variables, overcoming FBA’s linearity limitations.

Fig. S1 displays the sensitivity coefficients for the top-ranked parameters, showing first-order (direct impact) and total-order (overall contribution) sensitivity coefficients. A consistency among total-order coefficients suggests potential linear relationships between variables. This global analysis identifies parameters that need further research to enhance our understanding and reduce uncertainties. The input variable with the highest total-order sensitivity is tryptophan flux (*T rp*_*L*_), followed by teichoic acids (*DM*_*teich*_45_*BS*), pantothenate (*pnto*_*R*), and leucine (*Leu*_*L*). These findings highlight the importance of accurately defining flux boundaries for biologically relevant predictions.

SA results align with literature ((Karasawa *et al*., 1995)), indicating that cysteine, isoleucine, leucine, proline, tryptophan, and valine are essential for *C. difficile* growth. Indeed, the most sensitive input variable is *T rp*_*L*, followed by *Leu*_*L*. Dipeptides *Gly*_*Leu* and *Ala*_*Leu* also rank highly in sensitivity, underscoring their importance for bacterial growth. Teichoic acids and pantothenate, while sensitive inputs are less relevant for investigating host-pathogen interactions in CDI within our framework.

By ensuring that key metabolites and reactions are accurately represented and coordinated between dynamic and metabolic models, the harmonisation enhances the biological relevance and accuracy of unified models. This is particularly important for understanding complex systems like bacterial growth and metabolism, which are influenced by a large number of interacting factors.

Also, the improved integration and SA lead to more robust biological model predictions. This has direct implications for studying metabolic pathways, growth conditions, and potential interventions in bacterial systems such as *C. difficile*. While the harmonisation primarily focuses on metabolic and dynamic model integration, it also highlights the lesser relevance of certain inputs for investigating host-pathogen interactions in CDI. This specificity ensures that the research remains focused on the most impactful areas, enhancing the practical applicability of the findings.

##### Harmonisation formalism

We detailed information on essential amino acids (*Cys*_*L, Ile*_*L, Leu*_*L, P ro*_*L, T rp*_*L*, and *V al*_*L*) as ODEs, identified as the most influential input parameters. Using the GreatMod framework, we designed an Extended Stochastic Petri Net to facilitate information transfer between ODEs and the genome-scale metabolic network. This allowed the ODEs-based model to be fine-tuned to match the behaviour of the FBA model and experimental data. The harmonisation formal definition requires first the definition of a generic dynamic model with *n* interacting elements and *h* possible events (some of them might represent reactions) in terms of the ODEs system:

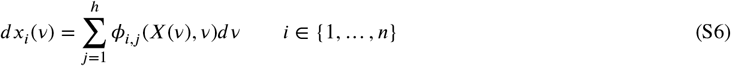

where *v* represents the time, *x*_*i*_(*v*) the number of the *i*-th element at time *v, X*(*v*) the vector of dimension *n* storing the number of each element at time *v*, and ϕ_*i*,*j*_ (*X*(*v*), *v*) ∈ ℛ the occurrence velocity of the *j*-th event considering the *i*-th element (if negative then the *i*-th element is consumed by the event, otherwise it is produced). Differently, a generic metabolic model with *m* metabolites and *r* reactions can be defined in terms of the FBA problem as in Eq.s S2.

Now the harmonisation between the two models is defined as follows. Defined *F* as the set of all the reactions in the FBA model, let *F* ^*bound*^ ⊂ *F* be the subset of the three types of boundary reactions: exchange, sink, and demand reactions. Thus, a set of boundary reactions exists, *F* ^*shared*^ ⊂ *F* ^*bound*^ , such that each corresponding flux can be associated with an occurrence velocity in the ODEs system, representing the exchange of such metabolites. We report in the Supplementary Section S1.3 more details about how to define *F* ^*shared*^ .

Mathematically, let *M*^*Ex*^ be the set of ODEs indexes representing the extracellular metabolites exchanged through the fluxes in *F* ^*shared*^ . We can define an objective function 𝒥 that maps an index from *M*^*Ex*^ into the index 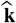 of the respective flux index, 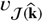. Indeed, if there exists an exchanged metabolite without an ODE associated, i.e., 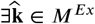 s.t.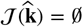, then a new ODE has to be defined to represent the concentration of the metabolite exchanged through the flux 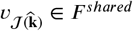. Thus, the ODEs associated with *M*^*Ex*^ will be characterised by a set of events that allows the connection between the starting ODEs model and the FBA model. So, the Eq. S6 becomes:

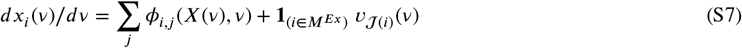

where 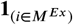 is the indicator function that returns 1 only if the index *i* ∈ *M*^*Ex*^. Furthermore, the fluxes are time-dependent since are conditioned to the FBA model which is continuously solved with respect to the state of the ODEs system at time *v*:

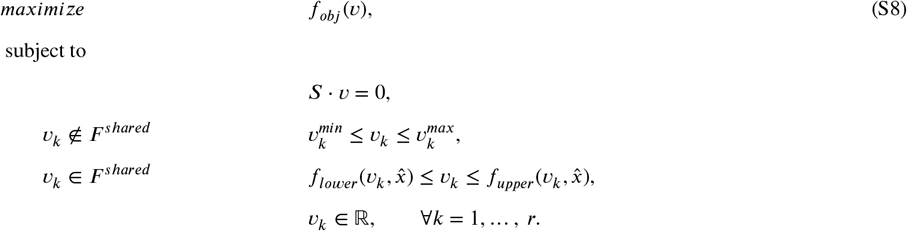

The functions *f*_*lower*_ and *f*_*upper*_ represent the lower and upper bounds in which each *v*_*k*_ flux varies, which can be defined as a linear transformation of the respective exchanged metabolites concentration (i.e.,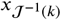) when *i* ∈ *M*^*Ex*^.In such a manner, the concentrations from the ODEs could be used as flux constraints because they represent the availability of the metabolites in the environment and therefore the uptake limit. Usually, the upper constraint is set only if a growth limit has to be defined, otherwise it should be infinity (see the Supplementary Section S1.4 for more details).

##### Harmonisation definition exploiting ESPNs as meta-formalism

The ESPN formalism becomes a meta-formalism that allows us to generalise and graphically harmonise the dynamic model with metabolic models by exploiting *general transitions*.Thus, given an FBA model as Eq. S2, we refine the set of general transitions, called *T*_*g*_ , into two disjoint subsets 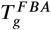 and 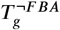, so that i) 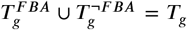, and ii) 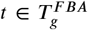 iff its intensity is obtained by solving the FBA model. Let us observe that, by definition, the transitions’ intensity has to be a positive number, therefore if reversible reactions are considered in *F* ^*shared*^ , then two different transitions, denoted with _*in* if the estimated flux is negative and _*out* if it is positive, have to be defined (an example is shown in Fig.1). Thus, we can infer that

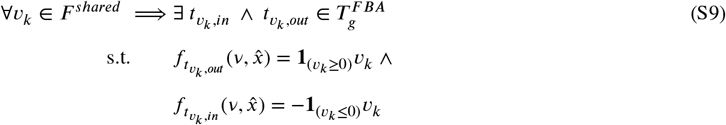

where, *f*_*t*_() is the intensity associated with the general transition *t*, 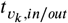 the general transition representing the flux *v*_*k*_, and **1**_(·)_ is the indicator function that returns 1 only if the input condition is satisfied.

Hence, the instantaneous changes of tokens *x*_*i*_(*v*) in the *i*^*th*^ place at time *v* expressed in the Eq. S7 can be rewritten as follows:

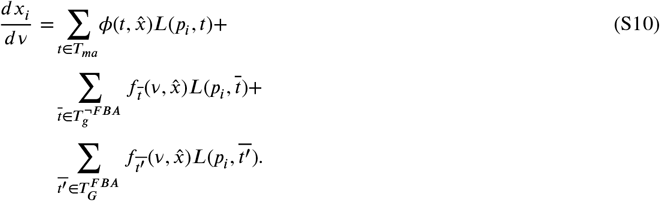

Thus, the 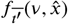 associated with a general transition 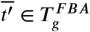 must be defined as in Eq. S9, with the fluxes *v* conditioned to the FBA model (Eq. S8) adding the dependence of the input places to the functions defining the bounds.

### S1.3 Reactions identification

Let us consider the FBA model defined in Eq. S2 in the main paper. Thus, we suggest following a few simple rules to identify the reactions which will characterize the set *F* ^*shared*^ as being efficiently coupled with the system of ODEs, as shown in Eq. S7 (main paper).

#### Considering boundary reactions

Firstly, we need to filter the set of all the fluxes of the FBA model, namely *F* , to consider the reactions that can be actually used to define *F* ^*shared*^ , i.e., the *boundary reactions*.Among all the reactions, we strongly suggest using only the boundary ones for two reasons: (i) to maintain consistency with the fundamentals of our paradigm, and (ii) not to violate the balance assumption of the FBA model. The paradigm has to connect two modelling approaches with their assumptions and rules, simulating two different system layers which share a given number of resources (e.g., metabolites). In particular, the FBA model is constrained by the steady state of the system, which supposes that there are no changes in any metabolite concentration in a specific time window. Therefore, by defining an ODE modelling the metabolite concentration over time of the FBA model, it is straightforward to see that the steady-state assumption of the FBA model does not hold anymore. Differently, the boundary reactions represent by definition the connection of the metabolic model with an extracellular environment, delineating in such a manner the perfect bridge between an ODE representing the shared metabolite outside the metabolic model, which acts as its resource by modifying dynamically the reaction constraints.

##### Sensibility of the model

Among the boundary reactions, the ones with a greater sensibility to perturbations in their constraints play a central role in the connection between the two modelling approaches. Given that the ODEs system is connected to the FBA model in terms of the fluxes constraints, which are defined by the functions *f*_*lower*_(·) and *f*_*upper*_(·), it is straightforward to consider reactions where small perturbations in the concentration of the shared metabolite affect the FBA solution. Otherwise, the two approaches could be solved independently without dynamic communication that might improve and change the analysis. Therefore, we suggest solving multiple times the FBA model varying the constraints of the boundary reactions to identify the ones that potentially will be the ones constituting *F* ^*shared*^ .

##### Reaction gap filling

Indeed, the knowledge of the system under study and the aim of the model plays a key role in the selection of these reactions, which can be exploited either to further filter, if just a few reactions are of interest, or to expand *F* ^*shared*^ with new reactions that are not present in the metabolic model. Considering the latter, the model could be either manually refined through the so-called gap-filling approaches. Reaction gap filling is a computational technique introduced in (Karp *et al*., 2018), by which it is possible to add a set of reactions to genome-scale metabolic models to obtain high-accuracy models. Gap filling completes what are otherwise incomplete models that are derived from annotated genomes in which not all enzymes have been identified. Thus, by adding specific boundary reactions to the FBA model, which will be part of *F* ^*shared*^ , it is possible to manually define exchanged metabolites that will connect the ODEs and FBA models.

### S1.4 Constraints definition

The functions *f*_*lower*_ and *f*_*upper*_ represent the lower and upper bounds in which each *v*_*i*_ flux varies, which may depend on the marking of the transition input places. Indeed, as reported in (Covert *et al*., 2008), several types of metabolic flux constraints can be used to define the *f*_*lower*_/*f*_*upper*_ functions:

1. **irreversibility constraints**, where the lower bound of the reaction is set to zero for reactions that can only proceed in the forward direction (Covert and Palsson, 2002);
2. **environmental constraints**, where the maximum flux through an exchange reaction is limited by the amount of substrate in the culture medium (Varma and Palsson, 1994);
3. **transport constraints**, which are represented as a maximum substrate uptake^2^;
4. **regulatory constraints**, where the flux through an enzyme is restricted by the expression of the corresponding protein(s) (W. Covert *et al*., 2001);
5. **ODE matching constraints**, where the fluxes are specified by the ODE model.

The first three types of constraints are derived from the literature (and it is the classical manner to define the constraints), so the functions can be expressed as 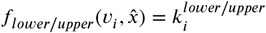, where 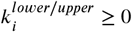 are constants. Indeed, when the flux does not depend on the ODEs values, then the 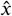 dependency can be omitted.

The regulatory constraints can be obtained by encoding in the function associated with the general transition a Boolean regulatory model, i.e., a set of Boolean logic equations that involve restricting expression of a transcription unit (sequence of nucleotides in DNA that codes for a single RNA molecule) to the value 1 if the transcription unit is transcribed and 0 if it is not. Similarly, the presence of an enzyme or regulatory protein, or certain conditions inside or outside of the cell, may be expressed as 1 if the enzyme, protein, or a certain condition is present and 0 if it is not. We decided not to consider this type of constraint but to focus on the ones that depend on the ODEs characterising the part of the ESPN which is not considered in the FBA. In this case, as shown in (Covert *et al*., 2008), the upper and lower bounds of the FBA fluxes depend on the entities modelled from the ODE system, and they are settled equally to the corresponding rate calculated by the ODE model. In such a manner, the concentrations from the ODEs could be used as flux constraints because they represent the availability of the metabolites in the environment and therefore represent the uptake limit. Usually, the upper constraint is set only if a growth limit has to be defined, otherwise, it should be infinity.

### S1.5 Synchronisation

Since we are implementing a hybrid model technique that combines two different algorithms for solving differential and algebraic systems, we have to define rules to synchronise these techniques. In particular, we define the ODEs system as the model leading the simulation, while the LPP characterising the FBA is exploited to calculate the value of some components of the ODEs system. Thus, according to this, we have to decide how many times and under which hypothesis the LPP has to be solved, and in this manner, the respective components in the ODEs system are updated.

In (Covert *et al*., 2008), the authors proposed a new type of DFBA, called integrated FBA, in which they integrated the FBA metabolic network with a Boolean transcriptional regulatory network and an ODE model. In particular, their simulations are characterised by a series of consecutive numerical integrations of the ODE model and solution of the LPP, whose interaction depends on some common metabolites and variables. In this context, they suggest choosing the length of each time step to be large enough that the FBA assumption (the concentrations of internal metabolites are time-invariant) holds, and yet small enough for the ODE model to calculate the system dynamics without accumulating numerical error.

Starting from this consideration, we defined a more general way to call the LPP during the ODEs solution without stopping it. In detail, we decided to solve the LPP only if the differences between all the input places of the general transitions modelling the FBA at time *t*_1_ and time *t*_2_, with 0 ≤ *t*_1_ < *t*_2_ < *t*_*final*_ (*t*_*final*_ is the final time selected for the ODE solution) is greater than ϵ > 0. Mathematically this can be expressed as follows (starting from the Eq.s S10 in the main paper):

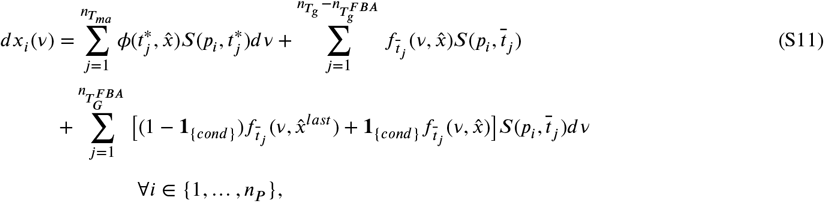

where **1**_{*cond*}_ is the indicator function which is 1 if the condition expressed in *cond* is true, otherwise is 0. Such condition can be defined as follows:

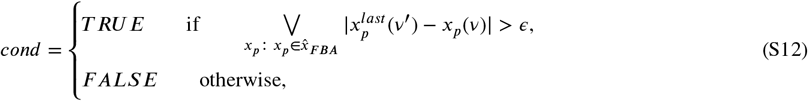

where 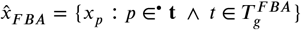 is the set of the markings of all the input places of the general transitions modelling the FBA, and 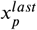 represents the marking of 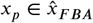 the last time *v*^′^ at which the FBA was calculated, with *v*^′^ < *v*. Therefore, the LPP is solved only if there exists a difference (defined by ϵ) between the only variables that could change the results of the FBA since the constraints of the flux associated with a specific general transition 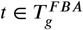 might depend on its input places marking. Otherwise, there would not be any important differences in the fluxes obtained from solving the FBA characterised by the same constraints (the only parameters of the FBA model that vary and depend on the ODE model).

### S2 *E. COLI* UNIFIED MODEL

An optimal test bed for our *UnifiedGreatMod*is given by the simulation of the carbon catabolite repression phenomenon exhibited by *E. coli* ((Ammar *et al*., 2018)). The bacterium adapts its metabolism to the environment’s available carbon sources (i.e., D-glucose or lactose). The LacZ system plays a crucial role in *E. coli*’s adaptive response to changes in available carbon sources. The role is to ensure that the cell expends energy on producing enzymes encoded by the lac operon only when necessary. To model this system at a genome-scale, we establish gene-protein-reaction (GPR) associations, connecting genes, proteins, and reactions via the flow of our novel modelling paradigm.

#### Overview of the modelling approach

In the iML1515 model, the lacZ gene encodes the enzyme beta-galactosidase, which catalyses the conversion of Lactose into D-glucose. The gene and the corresponding beta-galactosidase associated reaction are explicitly included (see Fig. S2D) in the metabolic network. By leveraging GPR associations, we employ Flux Balance Analysis (FBA) to predict metabolic fluxes. This allows us to simulate and study the activity of the LacZ system within the broader context of the metabolic network. Finally, integrating the regulation of the lacZ gene expression into our model provides a more accurate representation of the bacterial metabolic response to varying environmental conditions. For example, the model accounts for the fact that the lacZ gene is expressed only when lactose is present and D-glucose is absent.

Our approach combines experimental data, computational techniques, and environmental context to create a comprehensive computational translation considering a holistic view of biological systems. In this example, the unified model harmonises the diverse interacting domains of knowledge. We apply constraints derived from experiments to account for specific conditions. These constraints represent both intra- and extracellular factors, including (i) environmental restrictions, (ii) enzyme capabilities, and (iii) biomass maintenance needs. To implement these constraints, we set reaction flux bounds—defining lower and upper limits for each reaction. This ensures that our model adheres to the physiological boundaries observed in the biological system.

We develop a computational pipeline accounting for differential reaction activities and leveraging transcriptomics data. By mapping expression patterns of metabolic enzymes directly to changes in metabolic fluxes, we gain insights into the dynamic behaviour of the system. Specifically, we utilise data from *E. coli* obtained via the commercial microarray platform Affymetrix GeneChip system ((Liu *et al*., 2005)) as the expression data source. The expression data originates from MG1655 cells cultivated aerobically in MOPS minimal medium, supplemented with 0.1% glucose ((Bergholz *et al*., 2007)) representing the cell cultivation conditions. By tuning the bounds associated with exchange reactions, we simulate the availability of various nutrients in the environment. This integration allows us to capture both the internal metabolic state of the cell and the external conditions under which it grows.

#### Methodology

We used our computational workflow to construct the model of the metabolic system. The first step involved defining a mechanistic model and using Petri Net formalism to model the time dynamics of target metabolites. We then identified the metabolic network using the COBRA model of the *E. coli* K-12 strain MG1655. The next step was to identify the components of the metabolic model that would be coupled, which could include enzymes and metabolites involved in target metabolic reactions. After coupling the identified metabolic network, components, and mechanistic model, we converted metabolic reactions into an irreversible format to facilitate exchange reactions in Petri Net formalism. This involved splitting reversible reactions into two components and extending reaction identifiers with *in* or *out* to preserve original direction information.

The unified model, consisting of two places and six transitions, accounted for the metabolite concentrations required by the corresponding ODE model. The biochemical reactions of interest in the batch media were described by the Petri Net model, where transitions represented the inflow of fresh nutrients from the external environment. Finally, we used the coupled model to run simulations and predict system behaviour under various conditions. The model was used to demonstrate the direct effects of carbon supplementation on glucose and lactose consumption rates and bacterial biomass production in a biologically active environment. These rates were defined to model different carbon source administration regimes in a real-world batch culture scenario.

## Results

By following this approach, we gain insights into the dynamic behaviour of the LacZ system and its impact on overall metabolic processes. This study provides insights into the dynamics of the LacZ system and its influence on the metabolism of *E. coli*. We simulated different carbon source supplementation scenarios for a fed-batch. These include *constant feeding* (steady glucose and lactose rate), *linear feeding* (increasing glucose and lactose rate over time), *pulsed feeding 60* and *pulsed feeding 150* (adding glucose and lactose in short cycles), and *blank* (no glucose and lactose added). The prediction of sugar levels (Fig. S2A) shows a variation in exchange reaction activities (Fig. S2B) following the nutrient administration.

FBA resolution also estimates biomass flux (i.e. the cellular growth rate) under the given conditions (Fig. S2C). A biomass flux of about 0.65 mmol/gDW*h means that each gram of dry weight of the *E. coli* strain K-12 MG1655 is producing 0.65 mmol of biomass per hour. This value measures the metabolic activity estimated as the ability to convert the nutrients in the medium into new cell material. In a culture in the middle of its logarithmic growth phase, the estimated growth rate resembles the exponential growth phase (coli Genome Project University of Wisconsin–Madison, 2023). After the exponential growth phase, the cells have already utilised a substantial portion of the available nutrients, leading to a deceleration in their growth rate. *E. coli* efficiently uses glucose before its depletion due to its preferential metabolic pathway. The genome-scale metabolic network iML1515 includes a subnetwork for the uptake and metabolism of D-glucose and lactose (Fig. S2D), illustrating the dynamics of the lac operon. This subnetwork shows the transportation of D-glucose and lactose from the extracellular space to the periplasm and cytoplasm, where lactose can be cleaved into D-glucose by beta-galactosidase. The LacZ system, represented within this subnetwork, enables the cell to switch between D-glucose and lactose metabolism. This switch is governed by the availability of the sugars, with gene expression data and nutrient concentrations dynamically adjusting the bounds of key reactions (Fig. S2E).. Once glucose levels drop, the LacZ system is induced, enabling *E. coli* to use lactose as an alternative carbon source. The LacZ activity is constrained based on glucose availability to simulate the metabolic shift. When glucose is present, the LacZ system is repressed, but upon glucose depletion, the LacZ system is activated, allowing *E. coli* to use lactose after glucose depletion. This abrupt activation reflects the observed diauxic shift in *E. coli* metabolism. The switch on of lactose metabolism takes place around time 10 hours characterised by an increase of the bounds of reactions involving lactose uptake and beta-galactosidase activity when glucose levels are low and lactose is available and a fluxes rerouting through the subnetwork. This involves enhancing the transport of lactose from the extracellular space to the periplasm, the conversion of lactose to D-glucose in the periplasm, and the subsequent transport of D-glucose to the cytoplasm. (Fig. S2E, plot 2 and plot 4).

## S3 MODELING HOST AND DRUG RESPONSES TO CLOSTRIDIUM DIFFICILE INFECTION

### S3.1 Metabolic Basis of C. difficile pathogenesis

Intestinal epithelial cells ensure the transport of dietary amino acids through a process of uptake across the apical membrane, diffusion through the cytoplasm, and release via the basolateral membrane to the portal vein ((Argiles and Lopez-Soriano, 1990)). *C. difficile* is a minor member of the gut microbiota that often lives harmlessly in the small intestine so long as the ability of the gut microbiota to resist bacteria colonisation is maintained. However, some circumstances allow *C. difficile* overgrowth and disrupt the barrier function of the intestinal epithelium via enterotoxins production. The regulatory anti-starvation mechanisms are also involved in toxin production, corroborating that *C. difficile*’s pathogenesis is a blend between bacterial metabolism and the nutritional status of the environment.

The physiological effects of toxins involve mainly necrotic cell death increasing vascular permeability, affecting amino acid balance, and causing haemorrhaging. This might create a nutritional gut niche derived from the unleashing of nutrients from the host. There are some nutrients critical for efficient growth, which lack can trigger drivers of metabolic stress ((Neumann-Schaal *et al*., 2015)). Toxin-mediated inflammation alters the lumen metabolic pool of key amino acids (e.g. proline, leucine, isoleucine, valine, tryptophan and cysteine) establishing a direct link with gut environmental nutrient availability and to supporting increased bacterial growth (Edwards *et al*., 2014).

### S3.2 Physiological alterations during antibiotic perturbation

Guidelines about the awareness, diagnosis and treatment of CDI are available. However, several studies have reported treatment failure after an antibiotic course with varying extents of drug resistance (Debast *et al*., 2014). Given antimicrobial agents’ multifactorial action in different conditions, therapies informed by metabolic mechanisms of action will enable options with improved safety and efficacy. The metabolic alteration related to drug activation and deactivation is particularly suited to system-wide analysis that integrates multiple solution techniques. Therefore, we suggest a new computational approach to systematically investigate deregulated biochemical pathways on the landscape of metabolic changes concurrently with drug exposure.

The mechanism of action of different antimicrobial agents used to treat CDI is partly known. For example, nitroimidazoles require the reduction of the nitro group, which results in short-lived reduction products that oxidize DNA, followed by quick death in susceptible cells (Dingsdag and Hunter, 2017). The spectrum of activity depends on the selective action of the drugs on the specs of anaerobic metabolism. Nitroimidazole, changing into a component of the electron transfer system between pyruvate and hydrogenase, works as an electron siphon interrupting the electronic flow. The reduction product of the drug is directly responsible for getting cells killed. A sensitive organism undergoes several disrupted physiological mechanisms or effects together in a synergistic fashion.

### S3.3 Drug detoxification response modulated by micronutrients

We discuss novel findings regarding how micronutrients modulate the responses of both the host and *C. difficile* during infection. Various physiological conditions can alter micronutrient availability, contributing to infection susceptibility. Micronutrients necessary for regulating bacterial homeostasis may be derived from the inflammatory environment. Notably, micronutrients such as iron-coordinated heme, besides being indispensable for bacterial metabolism, function as redox-active molecules and redox stress defence factors (Knippel *et al*., 2018).

The inflamed gut contains numerous environmental oxidative stressors that can act as selective pressures, driving the evolution of mechanisms for oxidative stress protection. It is plausible that *C. difficile* acquires heme exogenously during infection, likely from the host due to toxin-mediated damage to the gastrointestinal epithelial layer, and endogenously from the diet. As an obligate anaerobe, *C. difficile* faces significant oxidative stress, particularly due to targeted antimicrobial therapy. The physiological effects of antibiotics used to treat anaerobic bacterial infections (e.g., nitroimidazoles) depend on the bacterial detoxification systems. This suggests that the acquisition of heme and the ability to withstand oxidative stress are crucial factors in the evolution of antibiotic resistance in *C. difficile*. These adaptive mechanisms enable the bacterium to survive in the hostile environment of the inflamed gut and resist antimicrobial therapy. Novel adaptive mechanisms may rely on disrupted metabolic pathways and reprogrammed bacterial metabolism, with necessary micronutrients for bacterial homeostasis derived from the inflammatory environment.

### S3.4 UnifiedGreatMod application: C. difficile infection

We combined the ODE-based dynamical model with the Genome-Scale Metabolic Model (GSMM) into a unified multiscale hybrid model. *C. difficile* strain CD196 GSMM was originally coded in the AGORA/MATLAB (1.03 version) format (Magnusdottir *et al*., 2016) and then elaborated using the R functions available at https://github.com/qBioTurin/epimod_FBAfunctions.git designed to integrate FBA-based model in R library *epimod*.

The metabolic model is parameterized using data of gut microbial metabolism and nutrition (Noronha *et al*., 2018). These nutritional data capture interactions in the context of human diet and microbiology. The coupling of ODEs and metabolic model dropped within one of the following scenarios: (i) when rates of ODEs and GSMMs reactions exactly match, the flux in the metabolic model was fixed to that in the dynamical model and represented as events using the ESPN description (ii) the subset of shared metabolites between both models are used for inbound communication in the metabolic model as constraints set. We presented a detailed example to illustrate the formalised representation method. The cellular component and molecular interactions involved in the CDI are defined by the model shown in (Fig. S3). From top to bottom, the model is organised into six modules: (i) *Blood vessel*), (ii) (*IECs*, (iii) *Lumen Env*, (iv) *C. difficile cells dynamics*, (v) *C. difficile metabolic network* and (vi) *Drug* modules.

We describe these modules in detail hereafter and report the mathematical information regarding the rates associated with each module’s transitions. By the definition of ESPN formalism, transitions that do not follow the Mass Action (MA) law are modelled as general transitions. In detail, the speed of general transition *t* ∈ *T*_*g*_ is defined by a function 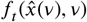, where 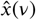 represents the vector of the average number of tokens for all the transition input places at time *v* and *t* is the transition. Differently, if *t* ∈ *T*_*ma*_, the velocity is defined as the MA law: a constant rate associated with the transition *t*, namely λ(*t*_*i*_), multiplied by the marking of all the input places of *t* powered by the cardinality of the arc, i.e., 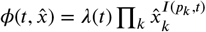. Let us observe that *x*_*place*_(*v*) denotes the average number of tokens in the place called *place* at time *v*.

### Blood vessel module

The transport of dietary amino acids into the bloodstream takes place across intestinal epithelial cells. This module can dynamically describe the transepithelial amino acids transported into the circulation following protein digestion and absorption. Most of the amino acids absorbed by epithelial cells (over 95%) are released in the bloodstream, whereas only a minor fraction is for cellular self-maintenance.

Transport transitions (referred to as *T*_*amino*_*e*_ remove ingested amino acids taken up from the gastrointestinal lumen and add the amount in the bloodstream. We gave these transition rates proportional to the concentration of metabolites that served as inputs to the transitions multiplied by a factor α (Table S1), assuming the nutrition amount restrained by epithelial cells.

### IECs module

The gut epithelium is the tissue comprising the intestinal epithelial cells. In this perspective, *IECs module* models intestinal epithelial cells dynamics and the absorption of nutrients by these cells. This module represents the processes related to altering the gut metabolic pool, such as the roles in nutrient absorption and death-induced nutrient release by IECs.

The state change refers to the transition *T*_*amino*_ ∈ {*T* _*pro*_*L*_*e, T* _*leu*_*L*_*e, T* _*ile*_*L*_*e, T* _*val*_*L*_*e, T* _*trp*_*L*_*e, T* _*cys*_*L*_*e*}, which removes at a constant rate from the system specific quantity of amino acids depending proportionally on correspondent input place *amino* ∈ {*pro*_*L*_*e, leu*_*L*_*e, ile*_*L*_*e, val*_*L*_*e, trp*_*L*_*e, cys*_*L*_*e*}. The speed of transitions is also proportionally related to the number of intestinal cells. These transitions are prevented from consuming the token from the *IECs* place, where each token represents a viable and functional IEC. The transition-associated functions are defined as:

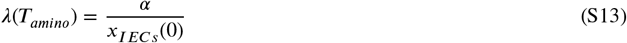

where α is the proportion of ingested amino acid taken up from the gastrointestinal lumen by enterocytes and transferred to the bloodstream. The term *x*_*IECs*_(0) is the initial number of cells forming the intestinal epithelium.

The initial number of cells was derived as follows. We consider that a single *mL* of cell culture (*C*_*IECs*_ equal to 2 * 10^5^ *cell*/*mL*) was plated in each Snapwell insert (well radius equal to 1.13 *cm*^2^) and cultured for two weeks, as described by (Anonye *et al*., 2019). The European Collection of Authenticated Cell Cultures (ECACC) reported that Caco-2 cells reach confluence after four days when seeded at 2 * 10^5^ *cell*/*cm*^2^ and since their doubling time is 84 *h*, we calculate that at the confluence, within a Snapwell insert, the number of IECs is 4.57 * 10^5^ cells.

Dietary heme is a crucial source of iron which is taken up into IECs and degraded by heme oxygenases. Then heme iron translocates across the cytosol and is finally released into circulation as the same pathway for non-heme iron. Heme iron absorption is far more efficient than non-heme iron absorption. *T*_*pheme*_*e*_ transition models the heme absorption. This function is defined as:

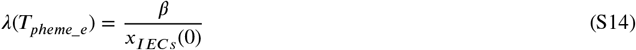

where *β* is the overall heme absorption yield. It is reasonable to expect that the overall dietary heme iron absorption yield is approximately 15–30% (Monsen, 1988).

The impact toxin harms on the mucosal intestinal response is displayed through its effects on epithelial cell death. *C. difficile* interaction with the single layer of intestinal epithelial cells leading to mucosal damage is modelled through the transition *IECsDeath*. This event represents IECs death. This transition is defined to represent the non-invasive pathogen features of *C. difficile* and the resultant toxin-mediated damage. Toxin production, which predominantly occurs during the stationary phase, is dependent on the bacterial amount. Therefore, *IECsDeath* velocity is proportional to the two input places *IECs* and *CD*:

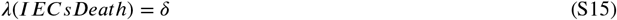

where *δ* is the number of host cells a single bacteria can kill. The initial marking for the place *Damage* equals to zero, indicating that simulations start with full membrane integrity.

Upon IECs death, a specific amount of amino acid is released in the lumen per dying cell. When IECs defoliate, amino acids deposited as protein are lost into the intestinal lumen. Therefore, we can evaluate the amount of amino acids released upon biomass wasting. For dividing cell types, the generic human biomass reaction available in human metabolic reconstruction is formulated taking biomass precursor as input and the biomass supermetabolite as output (Thiele *et al*., 2013). Biomass constraints are added to a biomass reaction by defining stoichiometric coefficients for each biomass precursor. Stoichiometry represents the intracellular amino acid quantity released in the extracellular environment upon each cell’s death. The arcs multiplicity was then computed given the amino acidic proportion constituting the prototypical human cellular biomass. Arcs connecting transition *IECsDeath* with output place *amino* is associated with multiplicity values detailing the number of tokens provided to target places:

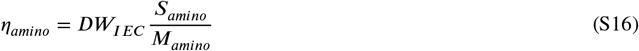

where *η*_*amino*_ represents the quantity (arc’s multiplicity) of amino acid injected in the output places *amino*. Each biomass unit lost several tokens, corresponding to the arc’s multiplicity. If an amino acid residual has molar mass (*M*_*amino*_), we can now convert the proportion into molar units.

### Lumen Env module

The gut lumen microenvironment is where the relations between bacterial colonizìsation, nutrient uptake and consumption occur. In the intestinal tract, this module models the toxin-mediated inflammation and the dietary intake of nutrients. This module allows for quantifying the gut metabolic pool value (i.e. essential amino acids and the micronutrient heme), which is the source of subsistence of bacterial growth during infection.

Minimal requirements of amino acids were previously determined for different strains (Karasawa *et al*., 1995). Cysteine, isoleucine, leucine, proline, tryptophan and valine were essential amino acids for the growth of *C. difficile*. Each amino acid is compartmentalised into environmental units, such as the extracellular space. The overlapping reactions between the ODE-based model and the metabolic model across this module are the exchange reaction to represent the intake (of efflux) of essential amino acid, *amino*, in the extracellular environment (see *C. difficile metabolic network* module).

The *Lumen Env* comprises 7 places and 11 transitions. The initial amount of amino acids was measured by (Adibi and Mercer, 1973) in the lumen of healthy human volunteers, except for tryptophan and cysteine whose concentrations were unknown. These values were inferred considering amino acid residual frequencies described in (Dyer, 2010), as explained below.

Aminoacidic composition is a type of basic feature of a protein sequence, which includes 20 discrete numbers, each representing the occurrence frequency of each of the 20 native amino acids in a protein sequence, respectively. Regarding the sum of all 20 encoded amino acids, the initial amount is given by the following relation:

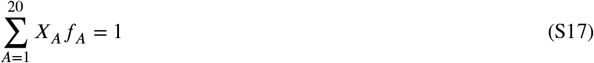

where *f*_*A*_ is the expected frequencies of amino acids, and *X*_*A*_ expected concentrations.

Transitions *D_pro_L_e, D_leu_L_e, D_ile_L_e, D_val_L_e, D_trp_L_e, D_cys_L_e* add to the system specific quantity of amino acid at a constant rate. Our model included the diet in the ODE-based model of the European Average diet. The nutritional data are collected at https://www.vmh.life (Noronha *et al*., 2018).

Consequently, nutritional flux units are then defined as:

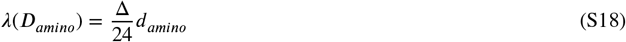

where λ(*D*_*amino*_) is the gross diet flux for amino acid residual *amino*, and *d*_*amino*_ is the nutritional flow for amino acid per person per day. We had to adapt available personal nutritional information to resize the nutritional flow to the model’s desired scale. For 1 *mL* of volume considered, we introduced Δ as metabolite flow scaling factor. The parameter’s value was set considering dietary flow *d*_*amino*_ at steady-state. We scaled to the amounts (*mmol*) as follows:

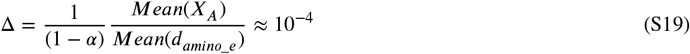

where α refers to the proportion of amino acid removed by the gut absorption activity. The factor Δ essentially converts the dietary intake rate from a per person scale (mmol/hperson) to a per mL scale (mmol/hmL), assuming that the concentration of amino acids in the gut is equal to the ratio of the concentration to the intake rate. This is a reasonable assumption under the assumption that the body is in a state where the intake and removal of amino acids are balanced, resulting in a stable concentration. The transition *D_pheme_e* models heme influx from nutrition is defined as:

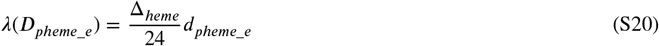

We chose to use heme as a significant model component as the inflammatory tissue damage causes the liberation of high concentrations of host heme at infection sites. The initial amount of heme, *x*_*pheme*_*e*_(0), was set as reported by (Hopp *et al*., 2020). Analogously, to scale the dietary intake rate from mmol/h per person to mmol/h per mL (the volume unit considered in our model), we introduced a conversion factor for heme, denoted as Δ_*heme*_.

Blood loss depends upon the integrity level of the IECs layer. Subsequently, red blood cells lyse once in the intestinal lumen, resulting in an abundance of extracellular heme, which is released from haemoglobin derived from ruptured erythrocytes under hemolytic conditions. Toxins cause severe damage to the colon’s intestinal epithelial layer, resulting in inflammation and bleeding. Free heme released from erythrocyte after intestinal haemorrhage causes an increase in intracellular free heme, this event is modelled by the transition *Inflam*.

IECs layer integrity could be monitored by measuring the transepithelial electrical resistance (TEER). The increase in host cell-associated bacteria was experimentally observed with a decrease in TEER by (Anonye *et al*., 2019), suggesting a disruption event of the intestinal epithelial barrier. The event *Inflam* represents several processes related to toxin-mediated epithelial integrity disruption. Since the elevation of inflammatory cytokines and chemokines, capillary diameter and blood flow increase allowing the extravasation of red blood cells. Consequently, the established inflammatory environment causes cytoskeleton rearrangement and membrane instability in red blood cells, which become more rigid and undergo lysis-releasing heme ((Gutierrez *et al*., 2021)). The heme-releasing reaction velocity modelled through the *Inf lam* general transition is defined by the following function:

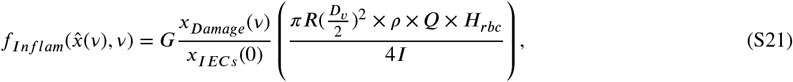

where *G* represents the grade of fenestration due to inflammation, *R* is the unbranched venules radius, *Q* is the red blood cell velocity, *ρ* erythrocytes density in human blood, *D*_*v*_ is the diameter of a villus blood vessel, and *H*_*rbc*_ is the amount of heme molecules within a erythrocyte. The villi of the small intestine project into the intestinal cavity, greatly increasing the surface area for food absorption and adding digestive secretions. Each villus is supplied with blood by 2 vessels, 1 venule, and 1 arteriole. *I* is the villus surface area, calculated as *I* = π × *W* × *L* × *m*, where *W* is the villus width and *L* is the villus height, and *m* is the surface amplification due to microvilli ((Solis *et al*., 2005)).

Bacteria have developed several strategies to capture heme and utilise the iron within the porphyrin ring. Utilisation of exogenous heme by bacteria involves the binding of heme to the cell surface receptors, followed by the transport of heme into cells. This reaction is mediated by the heme transport system represented by the transitions *HEMEti*_1_ and *HEMEti*_2_, whose speed is given by the following relation:

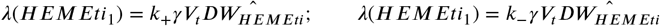

where *V*_*t*_ is the max velocity of the transport component that performs the uptake, *k* is the proportion of reversibility of heme transport reaction (+ for heme uptake; − for heme efflux), and γ is the proportion of mass protein which is heme transporter. 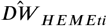 represents the average transporter amount of a *C. difficile* cell.

Subtraction of the endogenous uptake from the heme transport-induced uptake revealed a saturable component with an apparent *V*_*t*_ of 3.1 pmol/min/µ g protein ((Shayeghi *et al*., 2005)). Given the average protein content of a bacterial cell around 55% of the dry weight and the average molecular weight of proteins in the cell is around 50 kDa, we compute the total number of proteins in bacterial cells assuming that the number of transporter proteins is a small fraction of the total number of proteins. Considering that the transporter protein has a similar weight to the average protein, then the mass of the transporter protein in a unit of bacterial mass can be calculated as the ratio of the number of transporter proteins to the total number of proteins in the cell.

#### C. difficile cell dynamics mudule

This module models the proliferation response of a bacteria population upon treatment with antimicrobial therapy. This module is composed of 3 places: *CD, BiomassCD*, and *pheme_c*. This module also include 4 transitions: *DeathBac, Dup, Starv* and *Detox*. The presence of a token in the place *CD* is interpreted as a biologically active *C. difficile* cell. *C. difficile* initial cell number was calculated from (Lynch *et al*., 2013). Authors set the *in vitro* intestinal epithelial tissue as a planar IECs monolayer infected with a Multiplicity of Infection (MOI) of 100:1, *i*.*e*. 100 bacterial cells for each IEC. Given that we consider an initial number of IECs of 4.57 * 10^5^ cells, the initial number of *C. difficile* cells is 4.57 * 10^6^ cells.

The place *BiomassCD* indicates the average cellular biomass dry weight measured in *pg* per cell. *C. difficile* are Gram-positive rods, measuring 2-9 µ*m* in length (*a*) and 0.3-0.7 µ*m* in width (*d*) (Modaber, 1975). The cell shape shape can be approximated as a spherocylinder, a cylinder with hemispherical caps. Given the quoted diameter and length, we can compute a more refined estimate for the cellular biovolume of:

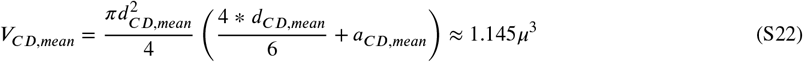

We can compute a more refined estimate for the biomass per cell of:

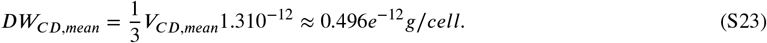

because most cells are about 2/3rd water, and the other components, like proteins, have a characteristic density of about 1.3 times the density of water.

The modelled system repurposes heme to counteract antimicrobial oxidative stress responses. The transition *Detox* the activity of hemoprotein HsmA to reduce oxidative damage caused by antibiotic action (Knippel *et al*., 2020). This event models the suggestion that the HsmA protein interacts with heme and the drug, and then returns to its original state, presumably after having altered the heme and/or drug in some way to reduce their toxicity or reactivity. This is consistent with recent findings which suggest that HsmA uses heme to protect *C. difficile* from oxidative stress. HsmA is a protein that contains a heme prosthetic group where the intracellular heme molecule could be the replaceable cofactor. The kinetic analysis of the system led to the scheme:

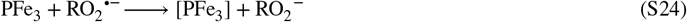

The rate constant for the reaction of radical reduction is equal to 1.5 * 10^10^ 1/mmol*min (Brault, 1985). The molarity of hemoprotein HsmA expressed by each bacterial cell is unknown. Therefore, the parameter *r*_*Detox*_ was defined by the model calibration phase.

The general transition *DeathBac* quantifies the clearance rate of bacterial cells by biomass minimum production. It is chosen according to the mean dry weight computed as shown in equation (S23). So the function is defined as:

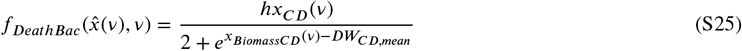

where *DW*_*CD*,*mean*_ represents one bacterial cell’s mean observed dry weight. The parameter *h* is the death rate representing lifespan decline due to depletion of nutritional resources, at which time the cell growth rate slows.

The difference between the average biomass of a cell in the bacterial population and the mean biomass for a bacterial cell observed in nature. It measures of how much the average biomass of a cell in the population deviates from what is typically observed in nature. This difference could have various implications depending on the specific biological context. For example, a positive value could indicate that the bacterial population is well-fed or experiencing favourable conditions, allowing them to grow larger than average. On the other hand, a negative value could indicate that the bacterial population is under stress or experiencing unfavourable conditions, causing them to be smaller than average.

The transition *Dup* models bacterial cell proliferation based on exponential biomass accumulation computed in each iteration until a threshold is reached and the cell divides. *Dup* is a general transition whose velocity is defined by the following function:

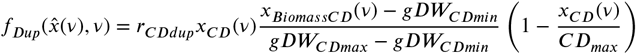

where *r*_*CDdup*_ is the doubling time, *CD*_*max*_ the theoretical maximum bacteria cell concentration, and *gDW*_*CDmax*_/*gDW*_*CDmin*_ the maximum/minimum biomass dry weight. We assumed that the maximum population for *C. difficile* at the end of the growth phase can reach a density of 10 billion cells per mL (Bressloff, 2014). *C. difficile* is a slow-growing bacterium. The generation time of *C. difficile* can range from 27 minutes to 216 minutes, depending on the strain and conditions. If we consider *r*_*CDdup*_ as the duplication rate per hour, a reasonable estimate might be between 0.28 (for a 216-minute generation time) and 2.22 (for a 27-minute generation time) divisions per hour.

The transition *Starv* models the recognition that microorganisms satisfy their maintenance homeostasis consuming biomass. Its parameter is defined as λ(*Starv*) = *EX*_*B*_*starv*, defined as the biomass flow at which the biomass is consumed for maintenance. A growth rate of 0.30 (1/h) requires biomass production in the range of 0.14 (mmol/gDW*h) (as a validation test, it is verisimilar that such biomass production requires glucose uptake of about 30 mmol/gDW*h)

### Metronizadole action mudule

This module models the mechanism of action of the antibiotic causing an imbalance between oxidative and antioxidative processes, inducing oxidative stress and cell death. This module is comprehensive of 1 place, *Drug*, which quantifies the amount of intracellular antibiotic. The module includes 3 transition: *Death4Treat, Efflux* and *Treat*.

The reduction of the drug promotes the formation of intermediate compounds and toxic free radicals. Free radicals interact with intracellular targets, especially with host cell DNA resulting in DNA strand breakage and deadly destabilisation of the DNA helix. Therefore, The efficacy of antibiotic treatment is dependent on the antibiotic amount to which the bacterium is exposed and on the intrinsic bacteria strain features which influence its susceptibility to therapy.

Transition *Death4Treat* represents the efficacy of antibiotic treatment at killing bacteria. Input places for this transition are *Drug* and *CD*. The multiplicity of the corresponding input arc connecting *CD* to *Death4Treat* quantifies the bacteria cell number lost for each unit of drug. The antimicrobial killing rate is defined as:

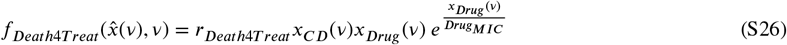

where *r*_*Death*4*Treat*_ represents the effectiveness of initial antibiotic therapy. The magnitude associated with this parameter was defined during the model calibration phase. The antibiotic concentration *Drug*_*MIC*_ is the Minimum Inhibitory Concentration (MIC) of the modelled antibiotic. The rate increases as the ratio of the drug concentration to its MIC increases. This could model the increased efficacy of the drug as its concentration surpasses the MIC.

The transition *Efflux* models the antibiotic efflux pathways that can be present in some bacteria strains, and it is defined by the constant parameter λ(*Efflux*) = *efflux*, where *efflux* represents the activity of Multi-Drug Efflux Transporter (MDET) that cause the efflux of intracellular antibiotics. We account kinetic parameters for the efflux pump from (Nagano and Nikaido, 2009). They cite a *V*_*max*_ of 2.35 * 10^11^ *mol*/*s*/10^9^ * *cell* which implies a total of 2.35 * 10^−12^ (*mol*/10^9^ * *cell* of transporter.

Fixing the frequency of daily dosing in the present study for simplicity, the transition *Treat* model the periodical antibiotic injections in the system. We designed the model to give an internal antibiotic concentration around the MIC, which implies a sub-prescribed *dose* of 0.05 (*g*/*day*). Every 8 h of time simulation the transition *Treat* adds an amount of antibiotic measured as *pmol* to the *Drug* place equal to 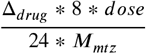, where *M*^*mtz*^ is the drug molecular weight (metronidazole), and Δ^*drug*^ is the scaling factor accounting for the amount of active drug reaching the site of action.

#### *C. difficile* metabolic network module

This module incorporates all metabolic reactions of the *C. difficile*. The metabolic level is connected with the *Lumen env module* by a set of 16 transitions. *EX_biomass_e_in* and *EX_biomass_e_out* coupled with the metabolic model exchange biomass reaction, and *EX_amino_L_e_in, EX_amino_L_e_out* coupled with the metabolic model exchange essential amino acid reactions. Transitions *sink_pheme_c_in* and *sink_pheme_c_out* represent connections of the reversible sink reaction for intracellular heme with the *pheme_c* place included in the *C. difficile cell dynamics mudule*. This reaction is functionally replaced by filling the undefined gap where heme is consumed by non-metabolic cellular processes but needs to be metabolized.

FBA identifies steady-state flux rates measured as *mmol*/(*g* * *h*) through a metabolic network, satisfying stoichiometric mass-balance as well as reaction direction’s constraints (the reaction’s flux activity represents the growth rate measured in 1/*h*, ((Adadi *et al*., 2012))). Metabolites are absorbed or released at an estimated rate and converted into biomass computed under the exponential growth phase. Biomass accumulation is limited to the maximal cell weight to restrict growth to physiologically feasible conditions. During optimisation, the upper bound of the objective function is set accordingly:

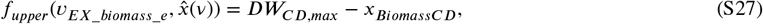

where *DW*_*CD*,*max*_ represents one bacterial cell’s maximum observed dry weight.

To incorporate flux balance analysis (FBA), we established constraints for the reactions *EX_cys_L(e), EX_trp_L(e), EX_val_L(e), EX_ile_L(e), EX_leu_L(e), EX_pro_L(e)*. The upper bounds for these reactions are set to zero, reflecting that there is no net production of aminoacids. The lower bound for each reaction, indicating the rate of metabolite consumption, is defined as follows:

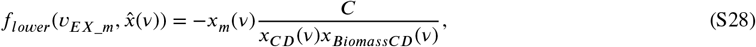

where *x*_*m*_(*v*) represents the concentration of the metabolite corresponding of the reaction *EX*_*m*, and C is a conversion scalar that bridges the units of concentration to the flux units in the FBA model. The lower bound in FBA, expressed in mmol/gDW/h, denotes the rate of metabolite consumption or production normalized to the biomass. A negative value for the lower bound indicates the consumption of the metabolite by the organism. To correctly map biomass values into the flux framework, considering that 1 pg (picogram) of biomass corresponds to 1 × 10^−12^ grams, we compute the conversion scalar *C* as follows: 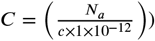, where *N a* is the number of molecules per mmol. Each token assigned to amino acids’ places counts *c* molecular units.

Considering the reaction *sink*_*pheme*(*c*), its upper bound is set equal to 10, and its lower bound is defined as follows:

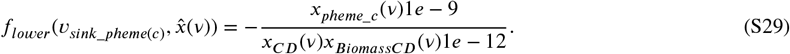

here, we need to correctly map flux values as *pmol* = 10^−9^*mmol*, analogously to S29. Finally, once the fluxes are estimated, the inverse functions of Eq.s S29 and S28 are applied to the correspondent *v* to convert it as a rate to associate with the transition.

### S3.5 Tables

**TABLE S2.**
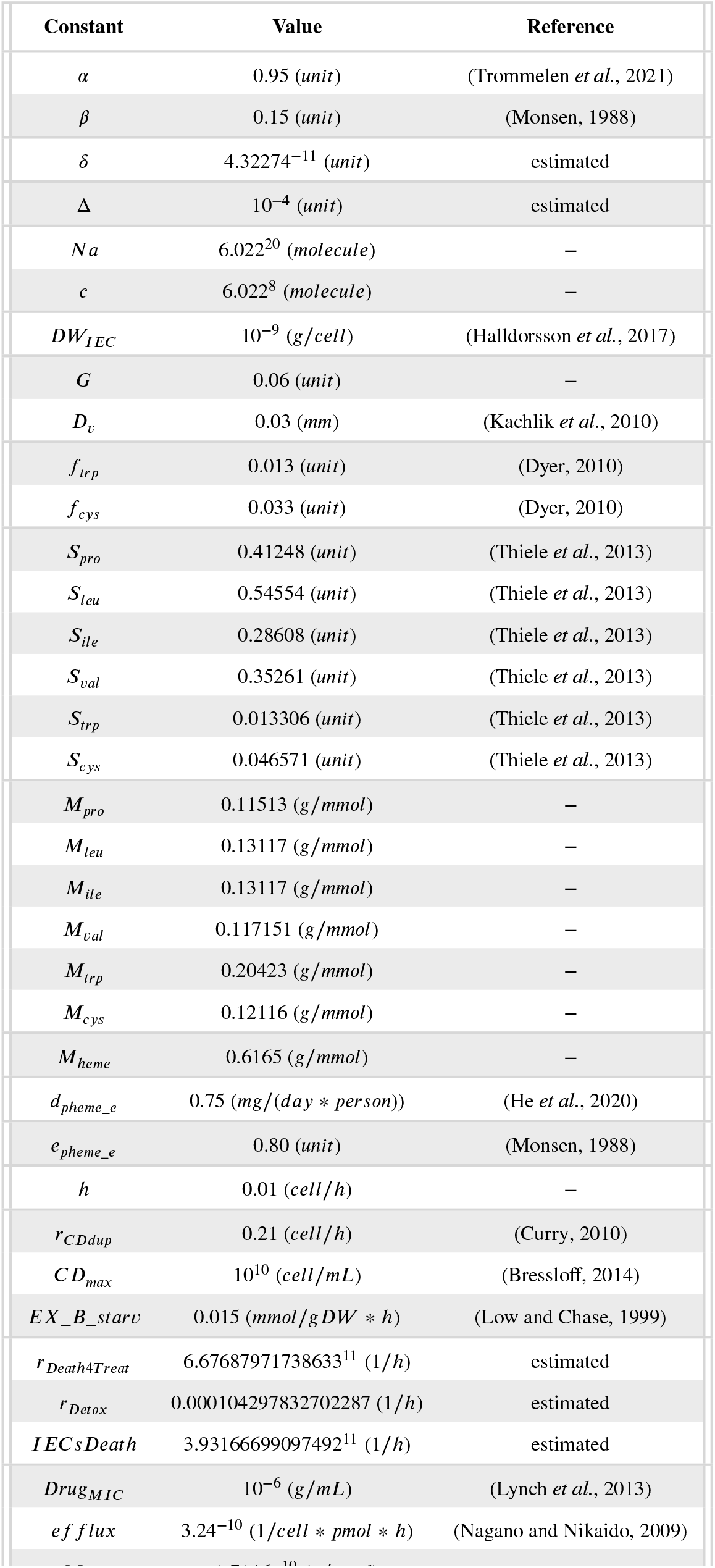
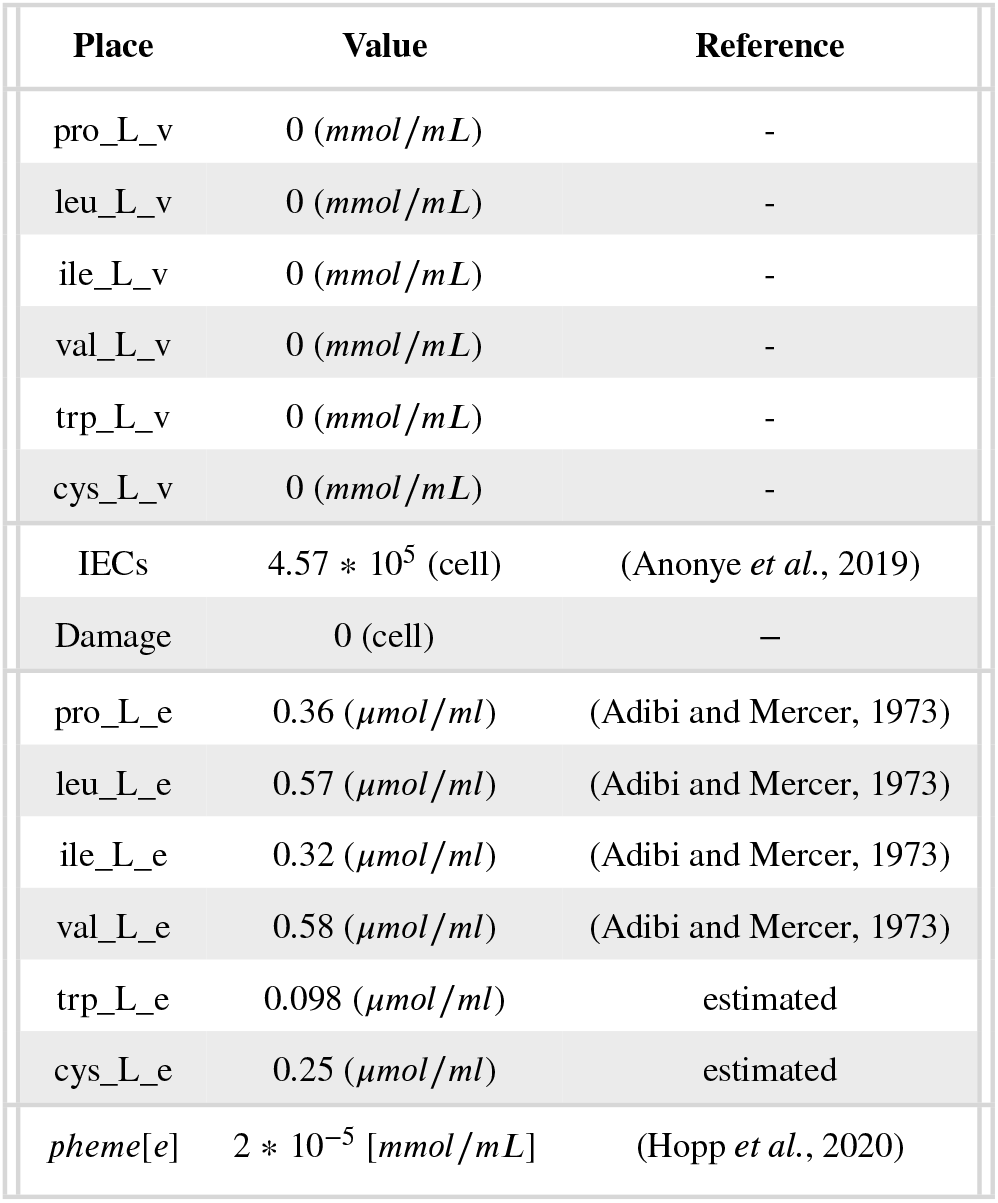
List of initial conditions of the dynamical model simulations.

### S3.6 Figures

**FIGURE S1.**
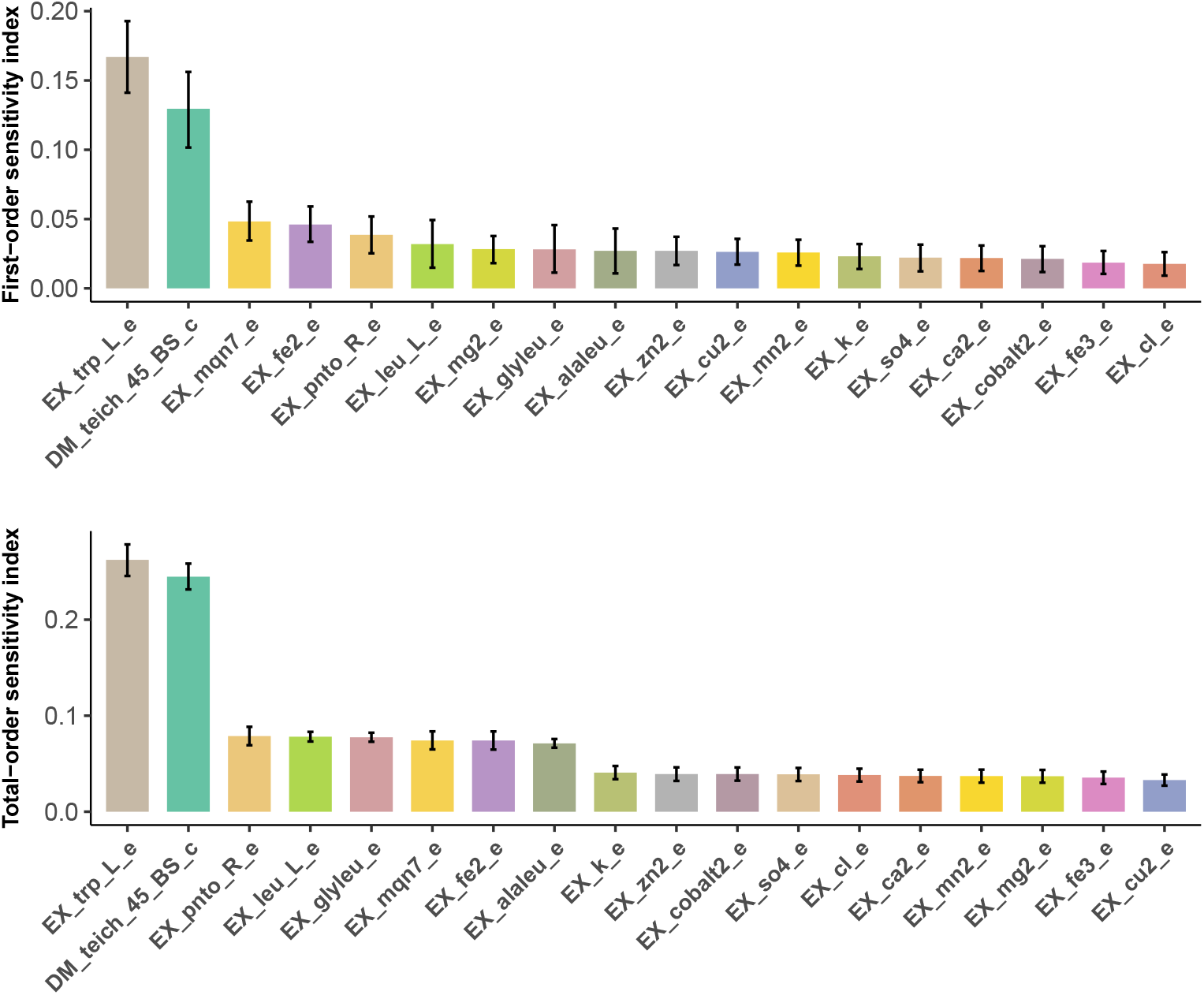
Results of the global SA on the *C. difficile* metabolic model. First-order sensitivity indices (top) and the total effect indices (bottom) and their low and high confidence levels. Bootstrapping is used to estimate the sampling distribution and to construct bootstrap confidence intervals on sensitivity indices. Sensitivity coefficients below 0.01 were considered negligible.

**FIGURE S2.**
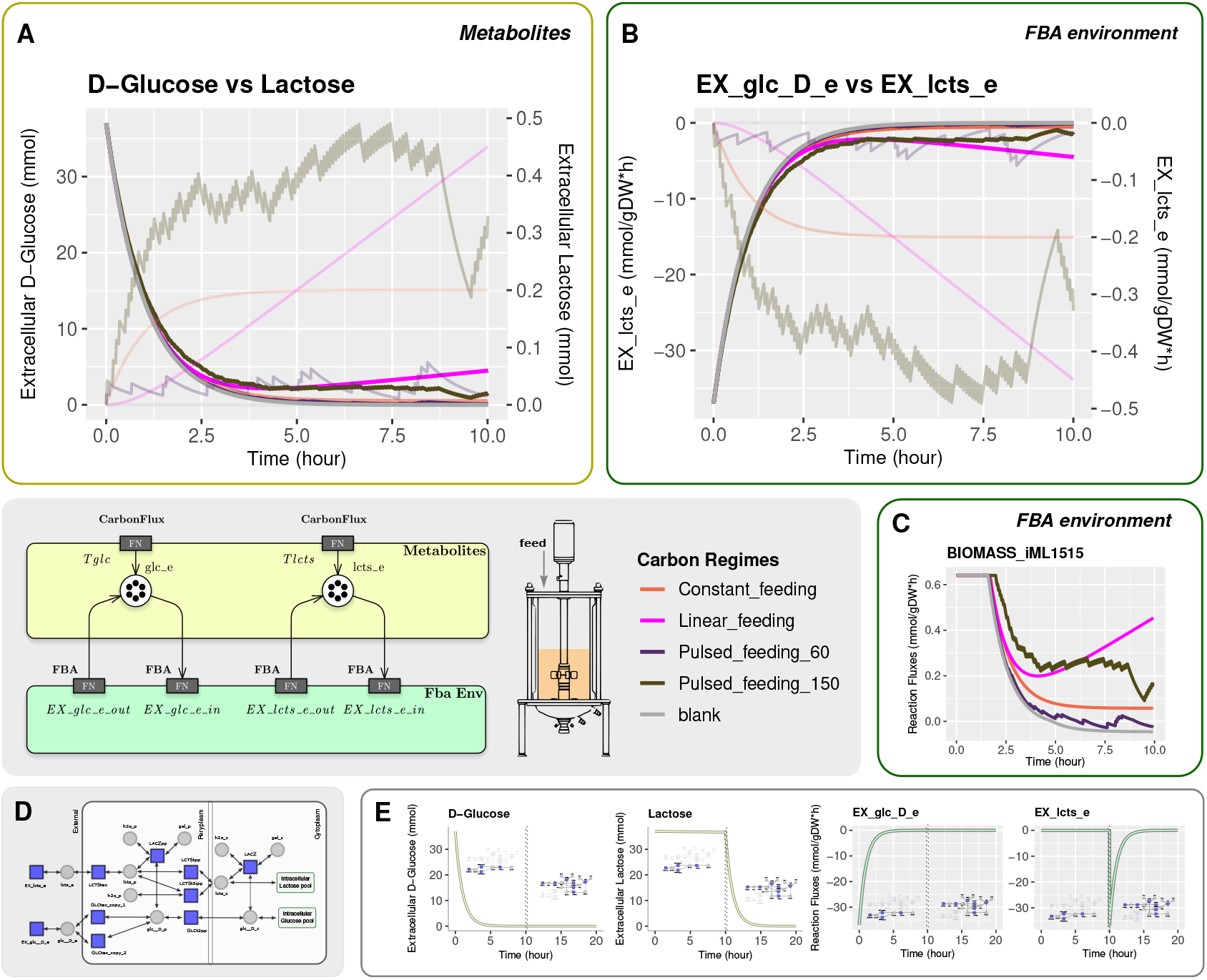
Overview of results for the *E. coli* batch growing model according to the supplementation scenario over time. (A) Profiles representing the time-course changes in the concentrations of D-glucose and lactose under different conditions of carbon administration. (B) FBA-estimated metabolic fluxes for D-glucose and lactose exchanges at time resolution. (grey panel) Petri Net model and schematic diagram of a bioreactor, (C) FBA-estimated *E. coli* biomass objective function at time resolution. (D) Graph of the subnetwork from iML1515 for the uptake of D-glucose and Lactose and components encoded by the lac operon. Grey circles represent the metabolites from the subnetwork, and blue squares represent the subsystem reactions. The pathway starts with the transport of D-Glucose and Lactose from the extracellular space to the periplasm, where lactose can be transformed into periplasmatic D-Glucose (glc_D_p) through a reaction from periplasmatic beta-galactosidase. D-glucose and lactose can also be directed to the cytoplasm where beta-galactosidase cleaves lactose into D-glucose. (E) Results collectively highlight the role of the LacZ system in enabling the switch from D-glucose to lactose metabolism when necessary.

**FIGURE S3.**
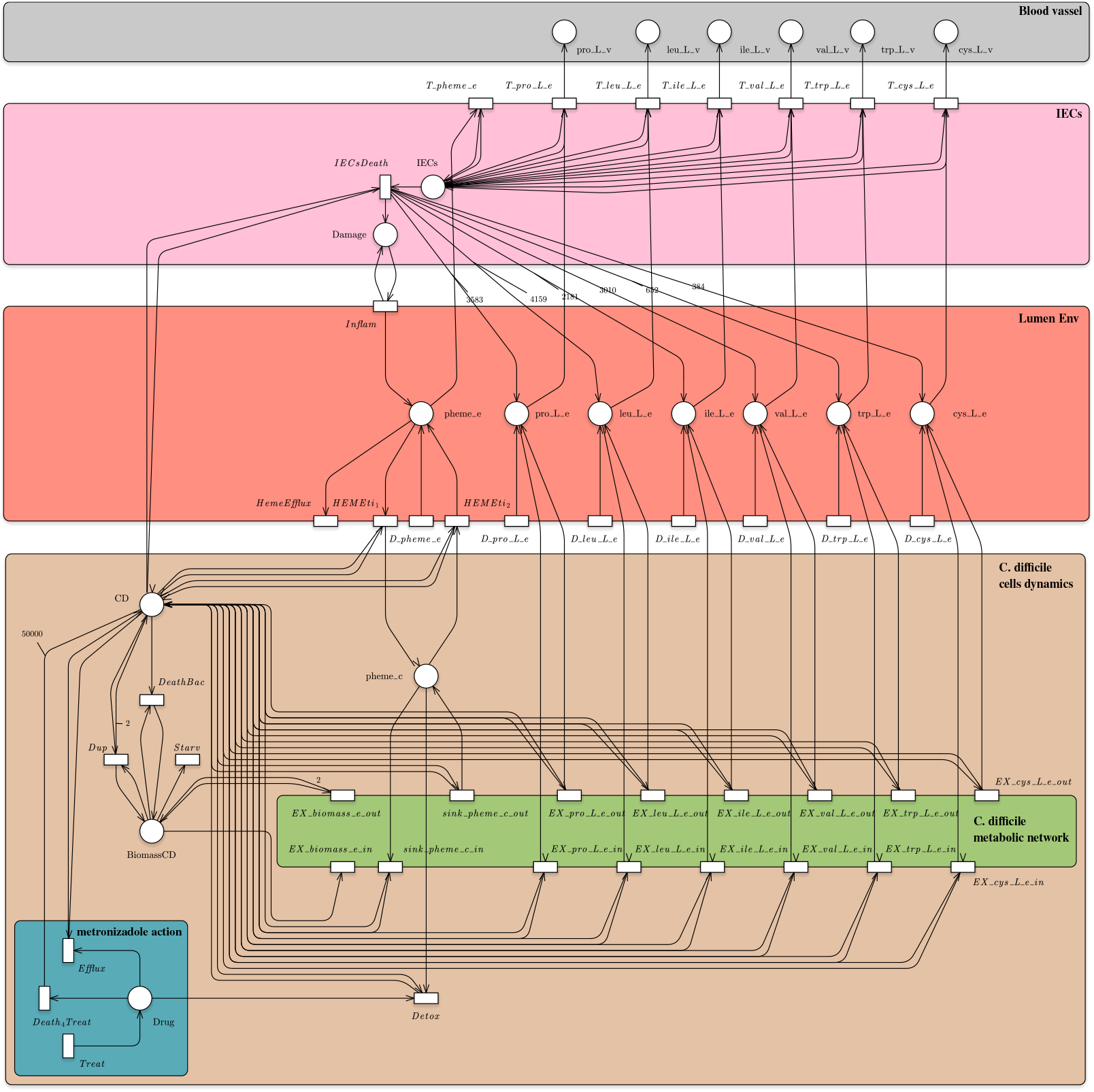
The ESPN associated with the CDI model is composed of places (graphically represented by circles) corresponding to epithelial cells, bacterial biomass, metabolites and tissue state (i.e. damage to the colonic mucosa), and of transitions (graphically represented by rectangles) representing interactions among the entities, cellular death, intake or efflux of metabolites, toxin action, intestinal inflammation and drug activity.

**FIGURE S4.**
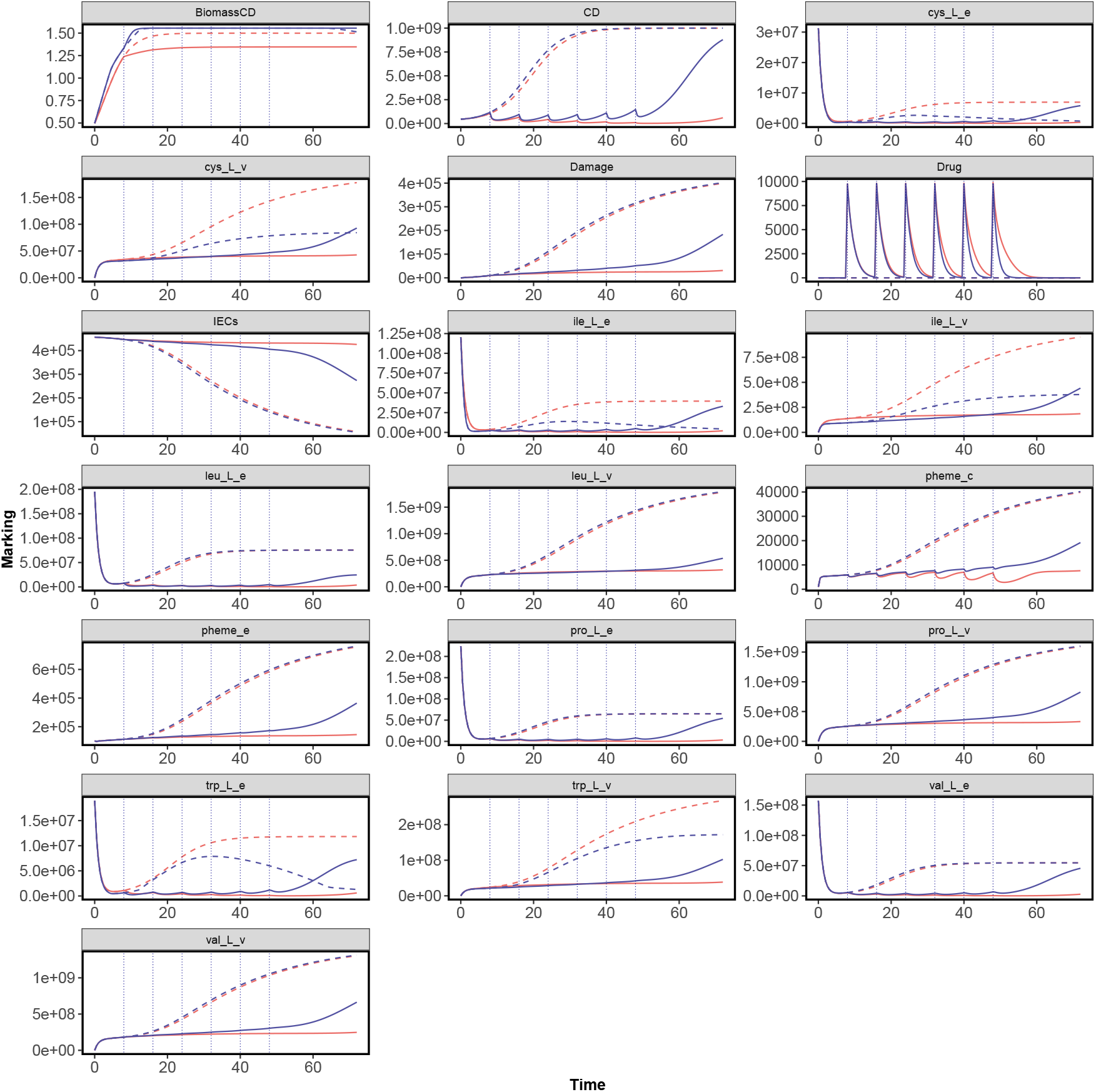
The dynamic behaviour of the principal places of the ESPN model given two scenarios, untreated and treatment conditions (red and blue colours, respectively), and the two approaches, unified and ablated (solid and dashed lines, respectively).

**FIGURE S5.**
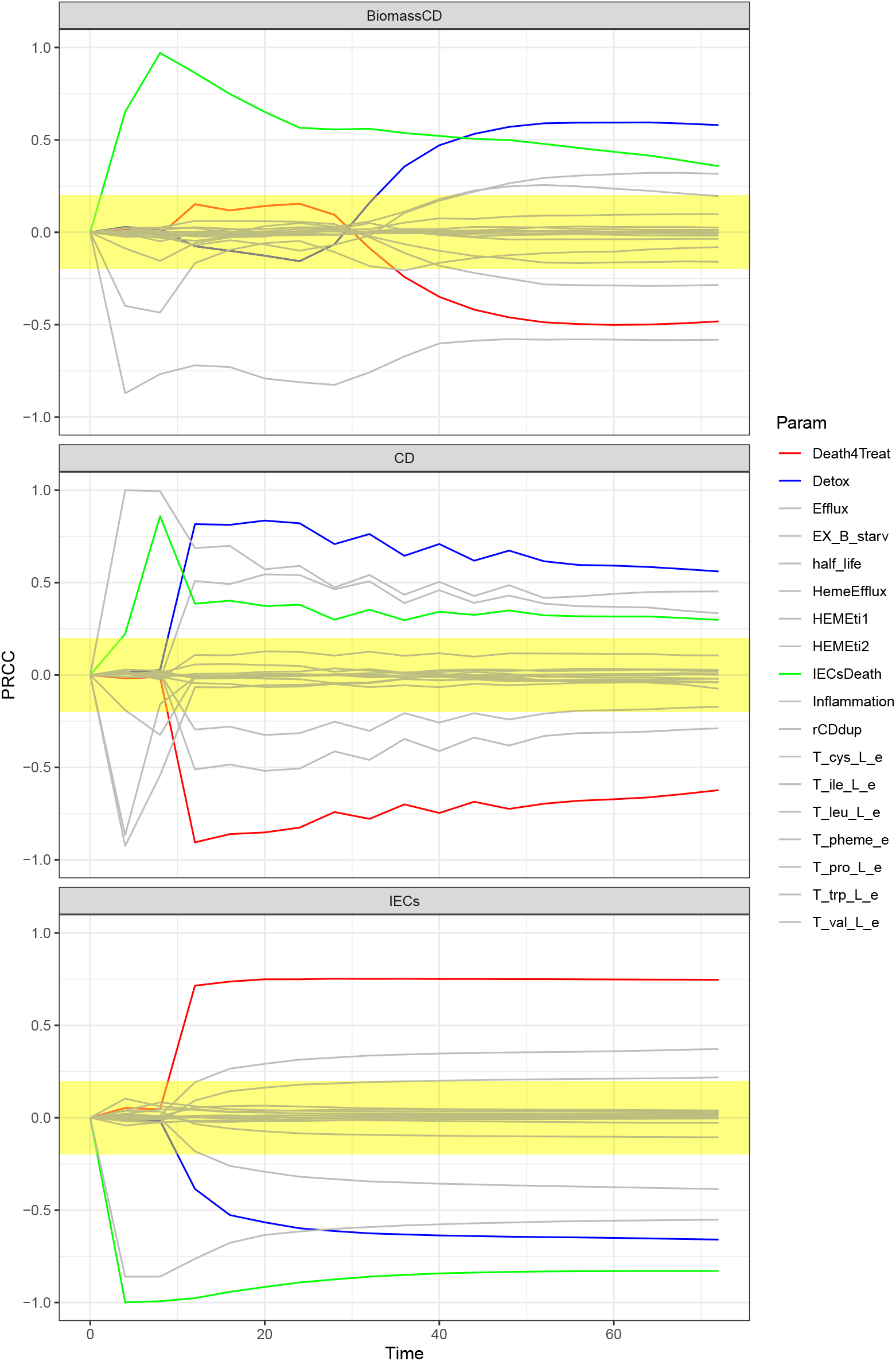
PRCCs over the whole time interval for each model parameter is reported. Yellow area represents the zone of non-significant PRCC values

## S4 ABLATION STUDY SETTING

To assess the significance of the unified framework, we proposed an ablation study only in the presence of drug injections. In context, we proposed to compare *UnifiedGreatMod* with two experimental designs: an *ablation experiment* and a *semi-ablation experiment*.

The first is obtained from the experimental design in which the dynamics of the metabolic environment are computed by FBA only at the beginning of the simulation. In the second experiment the FBA is computed at each time when an external stimulus occurs, i.e. at any drug injection. In both experiments, the flux estimated by the FBA considering the state of the model at that time is used as velocity for the associated transition. Given that the dynamic communication with the ODEs is lost, in order to take account of changes in the concentrations of the exchanged and shared metabolites outside the FBA model, the shared fluxes are normalized with respect to the concentration of the corresponding metabolite and then are used as constant rates in the Mass Action Law defining the velocity of the transition.

## Footnotes

1 The Backward Differentiation Formula method is implemented using the C++ LSODA library (https://en.smath.com/view/lsoda), while the optimisation problem using the *glpk* library (https://www.gnu.org/software/glpk/).

2 Typically the rate of uptake of nutrients is dictated by availability (a nutrient that is not present cannot be absorbed), concentration and diffusion constants (higher concentrations of quickly-diffusing metabolites are absorbed more quickly).

